# Microbiota-independent antiviral protection conferred by aminoglycoside antibiotics

**DOI:** 10.1101/248617

**Authors:** Smita Gopinath, Myoungjoo V. Kim, Tasfia Rakib, Patrick W. Wong, Michael van Zandt, Natasha A. Barry, Tsuneyasu Kaisho, Andrew L. Goodman, Akiko Iwasaki

**Author notes:** To whom correspondence should be addressed: Akiko Iwasaki, Ph.D., Department of Immunobiology, Yale University School of Medicine, 300 Cedar Street, New Haven, CT 06520.

## Abstract

Antibiotics are widely used to treat infections in humans. However, the impact of antibiotic use on host cells is understudied. We have identified a novel antiviral effect of commonly used aminoglycoside antibiotics. We show that mucosal application of aminoglycosides increased host resistance to a broad range of viral infections including herpes simplex viruses, influenza A virus and Zika virus. Aminoglycoside treatment also reduced viral replication in primary human cells. This antiviral activity was independent of the microbiota as aminoglycoside treatment protected germ-free mice. Microarray analysis uncovered a marked upregulation of transcripts for interferon-stimulated genes (ISGs) following aminoglycoside application. ISG induction was mediated by TLR3, and required TIR-domain-containing adapter-inducing interferon-β (TRIF), signaling adaptor, and interferon regulatory factors 3 (IRF3) and IRF7, transcription factors that promote ISG expression. XCR1+ dendritic cells, which uniquely express TLR3, were recruited to the vaginal mucosa upon aminoglycoside treatment and were required for ISG induction. These results highlight an unexpected ability of aminoglycoside antibiotics to confer broad antiviral resistance *in vivo*.

## Introduction

Antibiotics comprise a large and complex group of compounds typically secreted by bacteria targeting other prokaryotes. Antibiotics have revolutionized medicine since its discovery. However, overuse and misuse of antibiotics have led to the emergence of resistant bacteria in humans and in livestock. Current medicine faces a dire threat of antibiotic resistant bacteria, which is rampant in many hospitals around the world. In addition to its microbicidal activities, antibiotics can also directly affect mammalian cells in a variety of ways. Antibiotics have been reported to directly inhibit eukaryotic translation ^1^, inhibit mitochondrial function ^2,3^ and induce changes in mammalian metabolic pathways ^4^. There is a critical need to better understand the host effects of commonly used antibiotics.

Here, we found that topical delivery of an aminoglycoside antibiotic to the mucosa induced significant alteration of host gene expression, inducing increased expression of antiviral interferon-stimulated genes (ISGs). Aminoglycoside-mediated ISG induction resulted in significant protection against genital HSV-2 infection and significantly reduced Zika virus replication. Aminoglycoside-mediated antiviral protection was independent of the host microbiota, as intravaginal aminoglycoside treatment protected germ-free mice form HSV-2 infection. This protection mechanism operates in the vaginal mucosa and nasal mucosa, as intranasal aminoglycoside treatment increased ISG expression in the lung and aminoglycoside pre-treated hosts had increased survival after highly virulent influenza A virus infection. ISG expression was dependent on TLR3 and downstream signaling pathway, and robust recruitment of TLR3-expressing XCR1+ dendritic cells to the vaginal mucosa. Our results reveal an unexpected induction of antiviral state by commonly used aminoglycoside antibiotics.

## Results

### Prophylactic and post exposure application of vaginal antibiotics increase host resistance to genital HSV infection in a microbiota-independent manner

To investigate the effect of local, vaginal, antibiotic treatment on genital herpes virus infection, we treated mice daily with an antibiotic cocktail (composed of 0.5mg each of ampicillin, neomycin and vancomycin and 0.012mg metronidazole) administered intravaginally. All mice were first injected with Depo-provera to synchronize estrus cycles and increase HSV-2 infectivity^5^. After a week of daily antibiotic treatment, mice were infected intravaginally with HSV-2. After genital infection, HSV-2 undergoes multiple rounds of local replication in the vaginal mucosa which can be quantified by plaque assays for infectious virus in the vaginal wash ^6,7^. In mice, by four days post infection, the virus travels to the dorsal root ganglion where it can spread to fresh epithelial sites resulting in disease symptoms including hair loss and hind limb paralysis which can be quantified using a clinical scoring system^8^. Antibiotic-treated mice had significantly decreased vaginal viral titers and displayed fewer clinical symptoms of genital herpes infection indicating suppression of both early and late stages of HSV-2 infection (Fig. 1a,b). One day after infection, mice pre-treated with antibiotics had five fold lower viral titers in the vaginal mucosa with three out of the five antibiotic-pretreated mice having no detectable virus. By day 2, this difference increased to 500 fold indicating active suppression of viral replication in the mucosal epithelium (Fig. 1b). Vaginal viral titers of antibiotic-pre-treated mice eventually reached similar levels as those of control mice by days 3-6 post infection (Supplementary Fig. 1a-b). However, viral spread to the dorsal root ganglia (DRG) was significantly suppressed (Supplementary Fig. 1d). Neural spread of HSV-2 is required for disease as herpes viruses unable to replicate in neurons are unable to cause neurological symptoms and morbidity ^9^. In keeping with the reduced viral titers in the DRG, antibiotic pre-treated mice displayed little to no disease pathology as compared to control mice (Fig. 1a).

**Figure 1:**
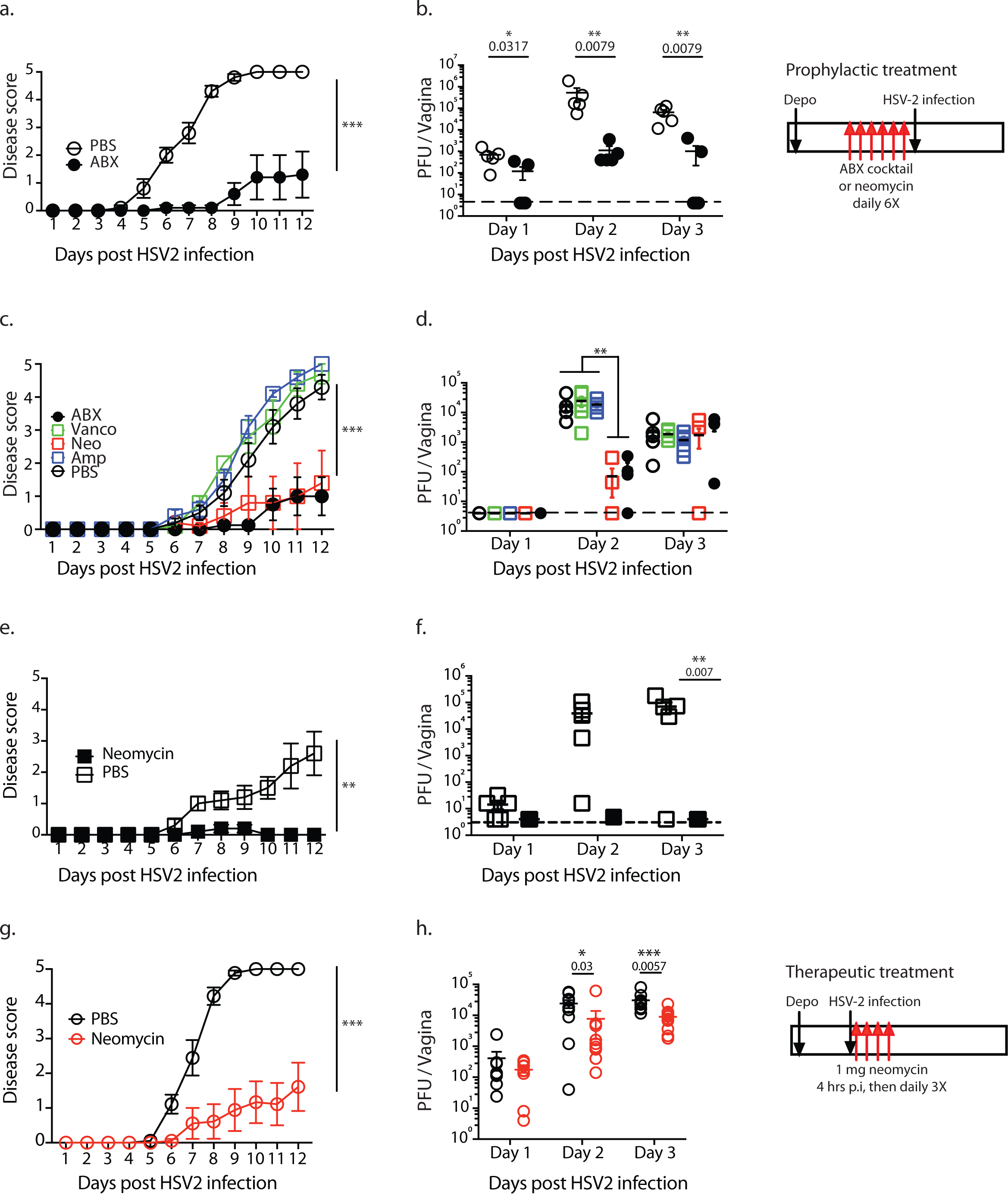
Vaginal application of neomycin confers prophylactic and therapeutic antiviral protection against HSV-2 in microbiota-independent manner. Conventional (a-d, g-h) and germ-free (e,f) mice were treated subcutaneously with Depo-Provera and five days after treatment were inoculated intravaginally with an antibiotic cocktail (a,b) or singly with the indicated antibiotic (c-h) daily for 6 days (n = 5 mice per group). After 1 week of treatment, all mice were infected intravaginally with HSV-2. For therapeutic neomycin treatment, depo-treated mice were infected with HSV-2, then treated with 1mg neomycin or PBS at 4hrs, 24hrs, 48hrs and 72hrs after infection (g,h). Disease score was monitored daily (a,c,e,g) and vaginal wash collected on the first three days (b,d,f,h). Error bars represent SEM. Significance was calculated using 2-way ANOVA. Exact p values are reported in Table S2.

To determine which of the antibiotics in the cocktail are required for protection against HSV infection, we treated mice singly with each antibiotic and compared disease progression to mice that had received the full antibiotic cocktail. We excluded metronidazole as it was present in very low quantities in the cocktail. Of the four antibiotics tested, neomycin alone recapitulated the host protection observed with the full antibiotic cocktail (Fig. 1c). Neomycin-treated mice had significantly lower vaginal viral titers as compared to other single antibiotic-treated mice and were identical to those treated with the full antibiotic cocktail (Fig. 1d). Thus, neomycin is the antiviral ingredient in the antibiotic cocktail.

Antibiotic-mediated effects on host immunity are often attributed to reduction or dysbiosis of relevant commensal bacterial communities. To determine if neomycin-mediated antiviral effect occurs via an impact on commensal vaginal flora, we treated germ-free mice intravaginally with neomycin. Notably, neomycin treatment of germ-free hosts also resulted in significant protection against genital HSV-2 infection. Neomycin-treated germ-free mice displayed no viral disease pathology (Fig. 1e) and had no detectable replicating virus in the vaginal mucosa (Fig. 1f) indicating that neomycin-mediated antiviral effect is independent of live or dead vaginal commensals, or microbiota in general. To determine if neomycin treatment had a therapeutic effect, HSV-2 infected mice were treated with neomycin at 4 hours post infection. Application of neomycin at later time points resulted in variable to no protection (data not shown). Neomycin treatment after infection resulted in lower viral titers (Fig. 1g) and highly reduced disease outcomes (Fig. 1h). These data indicated that vaginal application of neomycin before or shortly after exposure reduces HSV-2 infection and disease in a manner independent of commensal bacteria.

### Vaginal neomycin treatment induces expression of interferon-stimulated genes

To elucidate the mechanisms underlying neomycin-mediated host resistance against HSV-2, we analyzed vaginal gene transcription in neomycin-treated mice prior to infection. We identified over a hundred significantly upregulated genes (Fig. 2a; Supplementary Table 1). No genes were significantly downregulated in expression. Using Ingenuity pathway analysis, we found that genes in the type I interferon (IFN) pathway were heavily enriched (Fig. 2b), with over 30% of the upregulated genes falling within this pathway. We independently confirmed the upregulation of a subset of IFN-stimulated-genes (ISGs) using RT-qPCR (Fig. 2c). ISG expression was also increased in germ-free mice upon neomycin treatment (Supplementary Fig. 2). Neomycin-induced ISG expression was rapid, as a single treatment was sufficient to significantly increase ISG expression 2-5 fold though not to the levels observed after a week's treatment (8-10 fold) (Supplementary Fig. 3a). Neomycin-mediated ISG induction was also dose-dependent as increased amounts of neomycin correspondingly increased ISG expression (Supplementary Fig. 3a). Significant ISG induction was maintained up to 3 days after neomycin treatment and was only partially lost by 1 week post neomycin treatment with 5 of 8 genes still significantly upregulated (Supplementary Fig. 3c). Finally, this neomycin-mediated ISG induction was restricted to the site of application, as we observed no upregulation of ISG expression in the lungs of neomycin-treated mice (Supplementary Fig. 3d).

**Figure 2:**
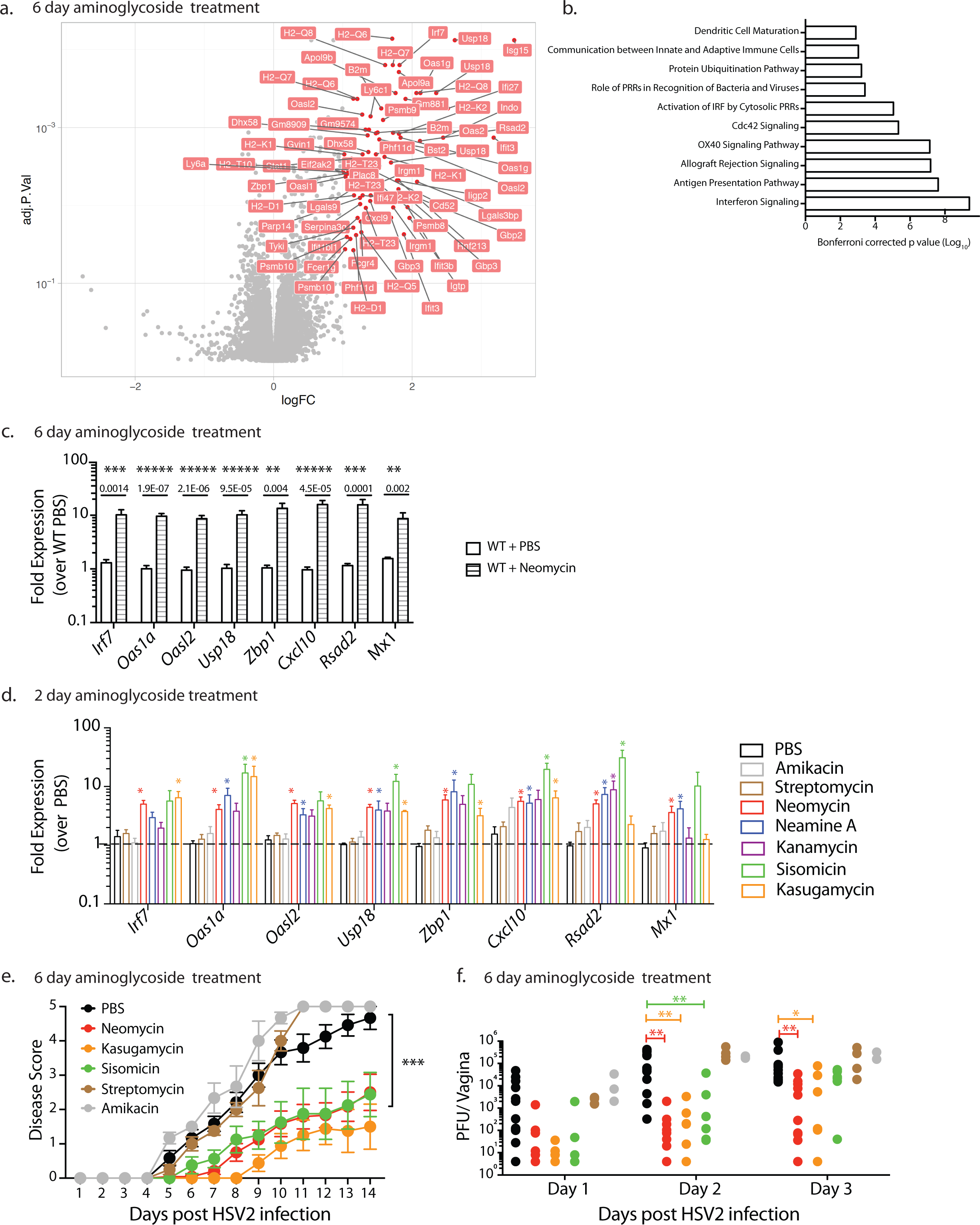
Vaginal application of most aminoglycosides induce interferon-stimulated genes, which is linked to antiviral protection. Microarray analysis of vaginal gene expression in neomycin-treated mice normalized against PBS-treated mice (n=3 per group) with fold expression and p values plotted (a). Significantly differentially expressed known genes (fold expression >1.5X, p value <0.05) are labeled. Ingenuity pathway analysis was used to identify the top ten signaling pathways enriched upon neomycin treatment (b). In an independent experiment Depo-treated mice were treated with neomycin or PBS (n=9) and vaginal gene expression measured via qPCR (c). Mice were treated subcutaneously with Depo-Provera and five days after treatment were inoculated intravaginally with the indicated aminoglycosides (1mg/day) for 2 days (n=3 mice per group). Vaginal gene expression was analyzed via qPCR and all comparisons were made against PBS samples (d). Depo-treated mice were treated intravaginally with 1 mg of the indicated aminoglycosides daily for 6 days and then infected with HSV-2 (e,f). Disease score was monitored (e) and vaginal viral titers measured (f). Error bars represent SEM and statistical significance was calculated using 2-way ANOVA. Exact p values for all comparisons are reported in Table S2.

### Most, but not all, aminoglycosides induce ISG expression

Neomycin is a member of a large and diverse group of aminoglycosides, many of which are commonly used to treat bacterial infection. To determine if ISG induction was common across the aminoglycoside family, we treated mice with a panel of structurally diverse aminoglycosides for 1-2 days. Five out of the seven aminoglycosides we tested significantly increased ISG expression upon application to the vaginal mucosa (Fig. 2d). The two non-inducers were streptomycin, which contains a streptamine core distinct from the other 2-deoxystreptamine-containing aminoglycosides, and amikacin, a kanamycin homolog that contains an addition L-haba side chain ^10^. Thus, the majority of the aminoglycosides tested induce ISG expression upon application to the vaginal mucosa. Since aminoglycosides mediate their antibiotic activity by binding ribosomal RNA, we tested if this ISG-induction was observed in other non-aminoglycoside ribosomal-targeting antibiotics ^11^. Mice were treated intravaginally with tetracycline or chloramphenicol, compounds that inhibit bacterial protein synthesis by binding the 50s ribosome subunit ^11^ (Supplementary Fig. 4a,b). Compared to neomycin, neither compounds induced high levels of ISGs, suggesting that ISG induction is not a common property of antibiotics that target the ribosome. Previous studies have reported increased mitochondrial dysfunction and a loss of mitochondria in mammalian cells ^2,12^ upon treatment with multiple classes of antibiotics. We observed no reduction in total mitochondrial DNA in neomycin-treated vaginal tissues (Supplementary Fig. 4c).

To determine the link between ISG expression and antiviral protection, we compared the ability of ISG-inducing and non-ISG inducing aminoglycosides to protect mice against HSV-2 infection. We selected two ISG-inducing aminoglycosides (kasugamycin, sisomicin) as well as the two non-ISG inducing aminoglycosides (amikacin, streptomycin). Mice pretreated with non-ISG inducers were not protected against HSV-2 infection, with vaginal viral titers and disease scores equivalent to those of PBS-treated control mice (Fig. 2e,f). Kasugamycin and sisomicin treatment however, resulted in significant reduction in vaginal viral titers and disease scores, displaying equivalent levels of antiviral protection as compared to neomycin (Fig. 2e,f). These data indicated that ISG induction by aminoglycosides corresponds to their ability to protect mice against viral challenge.

### Aminoglycosides induce antiviral protection against RNA and DNA viruses across multiple mucosal surfaces

Next, we examined if aminoglycoside-mediated induction of antiviral immunity was specific to the vaginal tract or if similar protection could be observed in other mucosal surfaces such as the respiratory tract. A single intranasal dose of neomycin sufficed to significantly upregulate ISG expression in the lung (Fig. 3a). To determine if this ISG induction results in host antiviral resistance, we tested neomycin-mediated protection in a functionally relevant mouse model of influenza infection. Many inbred mouse strains, including C57BL/6, lack Mx1 (Ref. ^13^). *Mx1* encodes the myxovirus resistance protein 1, a dynamin-like GTPase that blocks primary transcription of influenza by binding to viral nucleoproteins ^14-17^. Mx1 is an ISG that is induced by neomycin treatment (Fig. 3a), and when expressed,confers resistance to influenza A virus in mice^1414^. In mice lacking Mx1 such as C57BL/6 strains, innate resistance to influenza is abrogated, and they rely on adaptive immunity. We previously showed that oral neomycin treatment renders Mx1 deficient mice susceptible to influenza disease because it depletes gut commensal bacteria that normally support the function of dendritic cells that prime CD8 T cells^18^. In contrast, in this study, we used Mx1 congenic mice, which are highly resistant to influenza virus. The use of Mx1 congenic mice allowed us to study the ability of intranasal neomycin to elicit innate antiviral resistance, through inducing expression of ISGs including *Mx1*. To determine if neomycin-mediated ISG induction was sufficient to confer protection, we infected neomycin-pretreated Mx1 congenic mice with a highly virulent influenza A virus A/PR8 (hvPR8),which was selected for its ability to replicate rapidly even in the presence of Mx1 (Ref. ^19^). A single dose of neomycin was sufficient to significantly increase survival following challenge with an otherwise lethal dose of hvPR8, with 50% of mice protected (Fig. 3b). These results demonstrate that mucosal application of a single dose of neomycin can result in substantial protection against a highly virulent influenza A virus infection.

**Figure 3:**
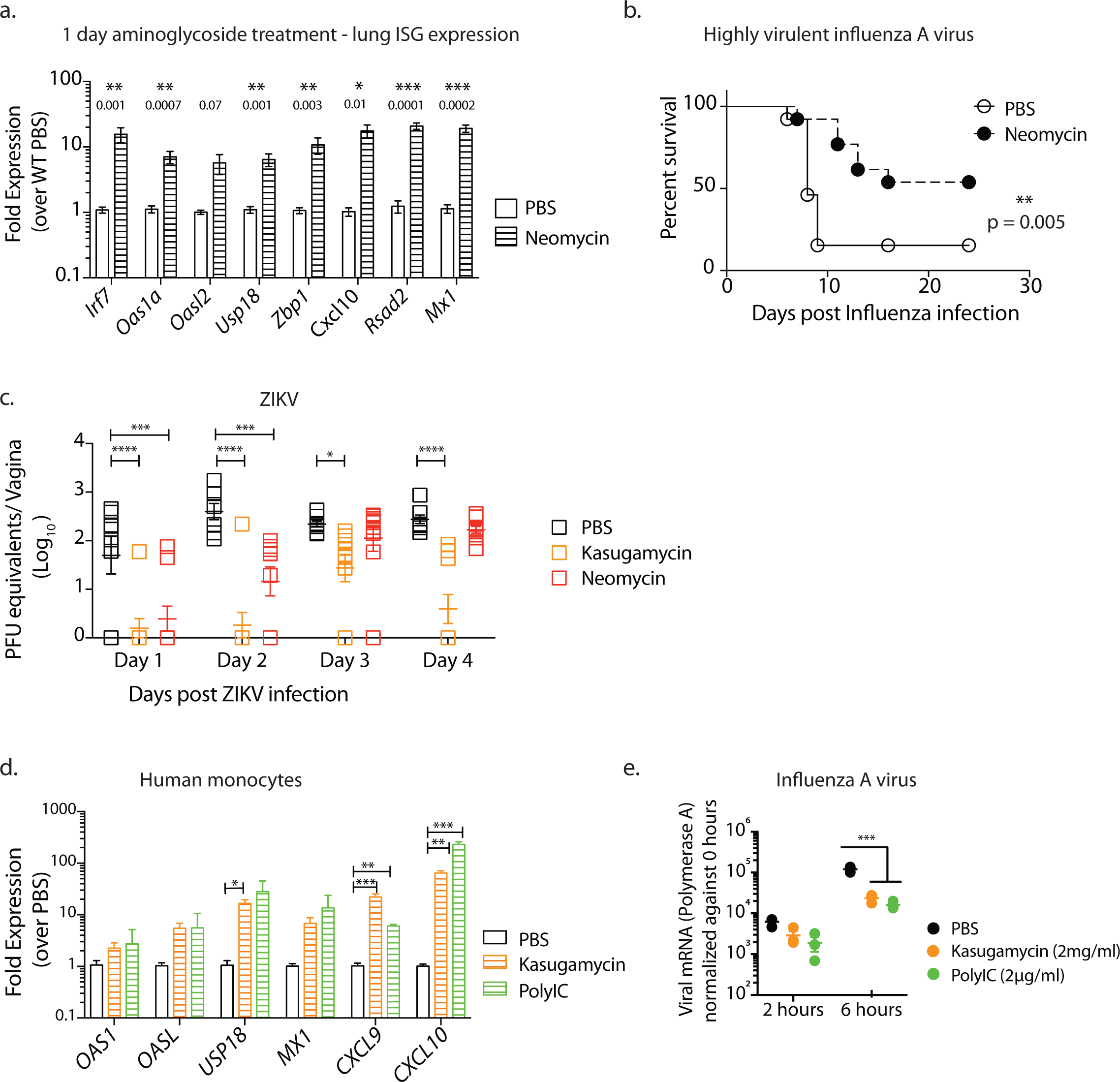
Aminoglycosides confer broad protection against both RNA and DNA viruses. Intranasal neomycin treatment induces ISG expression in lungs and protects mice against Influenza virus infection (a,b). Mice were treated intranasally with 2 mgs neomycin (n=5-7) and ISG expression analyzed 24 hours later in lung tissue (a). Mx1 congenic mice (n=7-8 per group) were pretreated with neomycin (2mgs) or PBS and infected 24 hours later with highly virulent influenza strain PR8 and survival curves compared using a log rank (Mantel-Cox) test (b). Depo-treated mice received intravaginal aminoglycoside or PBS daily for 6 days and were infected with 25,000 PFU ZIKV intravaginally and vaginal viral titers were calculated via qPCR (c). Primary human monocytes were treated with kasugamycin (2mg/ml) or Poly I:C μg/ml). Six hours after treatment, gene expression was analyzed (d) and in a separate experiment, kasugamycin-treated monocytes were infected with Influenza A/PR/8/34 (H1N1) strain at a multiplicity of infection of 2 and RNA collected 2 and 6 hours post infection. Virus levels were quantified via qPCR using primers against Polymerase A (e). Error bars represent SEM and significance was calculated using unpaired t-tests, correcting for multiple comparisons (a,d) or 2-way ANOVA (c,e). Exact p values for all comparisons are reported in Table S2.

We also tested the ability of aminoglycosides to protect against another RNA virus infection – Zika virus (ZIKV). ZIKV replicates in the vagina of wild-type mice and is controlled by IRF3 and IRF7-dependent type I IFN secretion and IFNAR signaling ^20^. Given the ability of aminoglycosides to induce ISGs, we examined if aminoglycoside-treated hosts are protected against ZIKV infection. We found that kasugamycin and neomycin treatment resulted in significantly lower levels of ZIKV RNA in the vaginal mucosa early in infection (Fig. 3c). Notably, while vaginal ZIKV titers in neomycin-treated mice eventually reached levels observed in PBS-treated controls, ZIKV replication in kasugamycin-treated mice remained significantly lower in a large proportion of mice in this group (Fig. 3c). To extend our findings to human cells, we treated primary peripheral blood monocytes obtained from healthy donors with kasugamycin. Kasugamycin treatment significantly increased ISG expression in human monocytes similar to Poly I:C treatment (Fig. 3d). Six hours of kasugamycin treatment was sufficient to induce robust ISG expression and significantly reduce replication of influenza A virus in monocytes (Fig. 3e). Collectively, our results demonstrate that aminoglycosides provide antiviral protection against a diverse set of DNA and RNA viruses: HSV-2, influenza A virus and ZIKV.

### Aminoglycoside induction of ISGs is independent of cytosolic DNA sensor signaling

Since neomycin treatment resulted in increased expression of genes in nucleic acid sensing pathways, we asked if cytosolic or endosomal nucleic acid sensing pathways were involved. To test the requirement of cytosolic nucleic acid sensors, mice lacking the RIG-I like receptor signaling adaptor *Mavs*, or the cytosolic DNA sensor cGAS, and its downstream adapter STING (*Tmem173*) were treated with neomycin or PBS and subsequent vaginal gene expression was examined. While neomycin induction of ISG expression was intact in cGAS or STING knockout mice, neomycin-treated MAVS knockout hosts showed relatively lower levels of ISG induction (Supplementary Fig. 5a,d). Infection of neomycin-treated MAVS knockout hosts resulted in reduced protection (Supplementary Fig. 5b,c). By contrast, neomycin-treatment of STING knockout mice resulted in robust antiviral protection with significantly suppressed viral replication and little to no disease pathology (Supplementary Fig. 5e,f). These results indicate that the cytosolic DNA sensor signaling pathway is dispensable for ISG induction by neomycin, while RNA sensor signaling pathways play a partial role in aminoglycoside-mediated ISG induction.

### Neomycin mediates antiviral effect by inducing ISG expression via activation of the TLR3-TRIF-IRF3/7 pathway

We next investigated the role of endosomal RNA sensors. While TLR7 signaling was dispensable for aminoglycoside-mediated antiviral protection (Supplementary Fig. 6a,b), we found that treatment of *Tlr3*^-/-^ mice with neomycin resulted in no ISG induction (Fig. 4a) and this was accompanied by significant loss of protection against HSV-2 infection with higher disease score and vaginal viral titers (Fig. 4b,c). TLR3 signals via the adaptor protein TRIF^21^. To confirm the activation of TLR3 signaling pathway in aminoglycoside-mediated antiviral protection, we treated *Trif*^-/-^ mice with neomycin. ISG induction was diminished in *Trif*^-/-^ mice as compared to neomycin-treated WT mice (Fig. 4d). This lack of ISG induction was accompanied by a lack of protection against genital HSV-2 infection, as neomycin-treated *Trif*^-/-^ mice had equivalent vaginal viral titers and similar disease scores as compared to PBS controls (Fig. 4e,f). Finally, we investigated the role of transcription factors downstream of TLR3 and TRIF signaling. ISGs downstream of cytosolic and endosomal nucleic acid sensors require IRF3 and IRF7 for induction. *Irf3*^-/-^ mice were treated with neomycin or PBS for one week and vaginal gene transcription analyzed. Similar to *Trif7*^-/-^ and *Tlr3*^-/-^ mice, no upregulation of ISGs was observed in neomycin-treated *Irf3*^-/-^ *Irf7*^-/-^ mice as compared to PBS-treated controls (Fig. 4g) and this lack of ISG induction resulted in lack of protection against HSV-2 infection (Fig. 4h,i). Induction of ISGs via IRF3 and IRF7 often requires signaling through the type I IFN receptor (IFNAR) ^23^ but IFN-independent ISG induction has also been reported ^24^. We observed increased induction of ISG expression upon neomycin treatment in *Ifnar1*^-/-^ mice suggesting that IFNAR signaling is dispensable (Supplementary Fig. 7a). However, since basal ISG expression was much lower in *IfnarT*^-/-^ mice as compared to WT, neomycin treatment only increased gene expression to those of untreated WT mice, thus neomycin treatment was not accompanied by significant antiviral protection (Supplementary Fig. 7b,c). Collectively, our results suggest that neomycin induces ISG expression via activation of TLR3-TRIF-IRF3/7 signaling pathway to confer protection against HSV-2.

**Figure 4:**
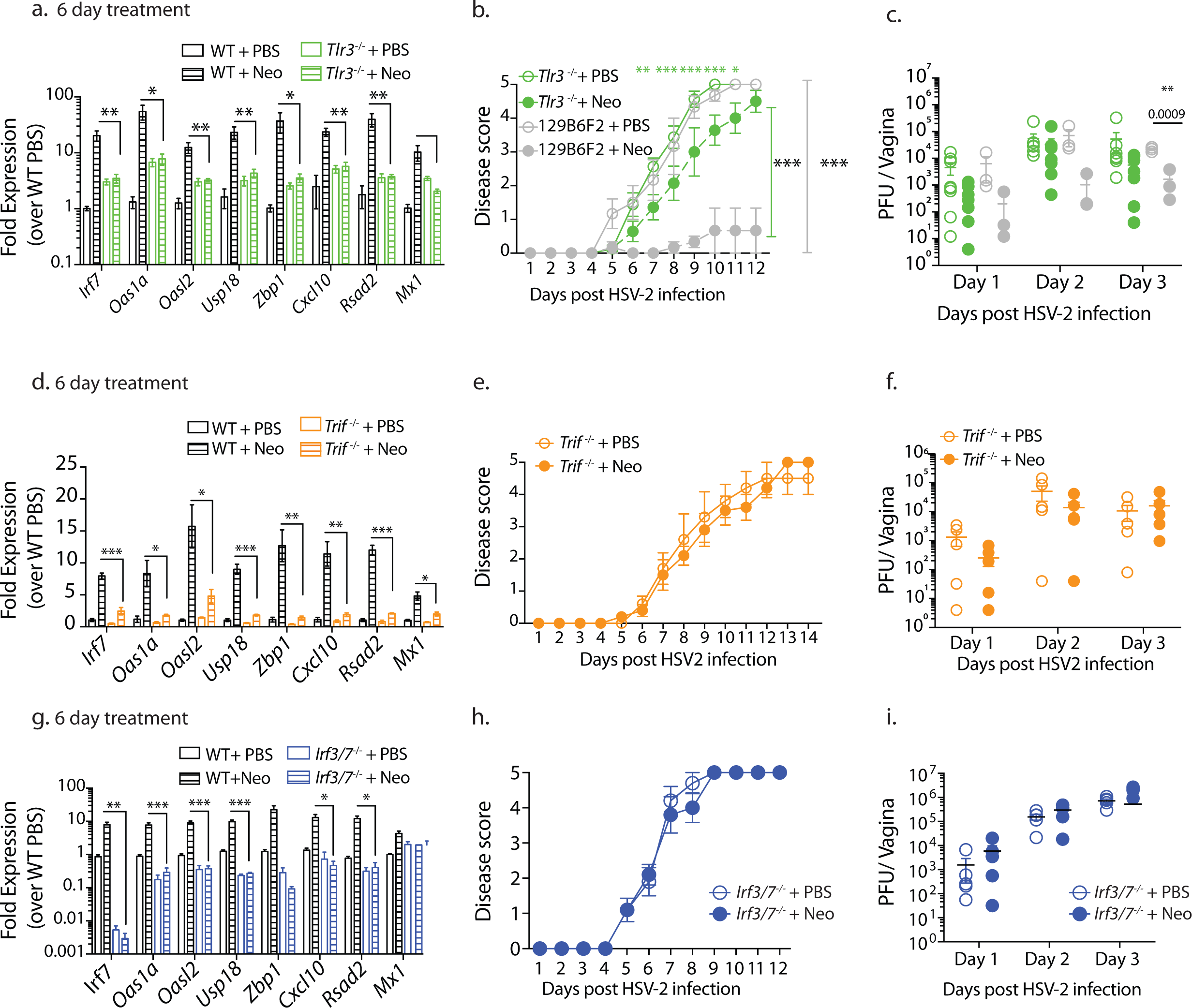
Aminoglycosides mediate antiviral immunity via the TLR3-TRIF-IRF3/7 signaling pathway. Depo-treated wildtype and knockout mice of the indicated genotypes (n=3-5) were treated with 1mg neomycin daily for 6 days and vaginal gene expression measured (a,d,g). In an independent experiment these mice were infected with HSV-2, disease scores (b,e,h) and vaginal viral titers measured (c,f,i). 129S1×B6 F2 mice were used as wild type controls for TLR3^-/-^ mice (b,c) and C57BL/6N mice were used as wildtype controls for remaining (a,d-i). Error bars represent SEM. Significance was calculated using 2-way ANOVA (b,e,h) or unpaired t-tests correcting for multiple comparisons. Exact p values for all comparisons are reported in Table S2.

### Recruited dendritic cells are required to induce ISGs following neomycin application

We next examined the cell types responsible for ISG induction and antiviral protection in the vaginal mucosa upon neomycin treatment. Since the vaginal mucosa is composed of stratified squamous epithelial cells, small numbers of resident leukocytes and circulating leukocytes, we investigated whether ISG induction by neomycin requires tissue resident cells or circulating leukocytes. To determine if circulating leukocytes were responsible for increased ISG expression, we blocked cellular recruitment by treating mice intravaginally with pertussis toxin. Pertussis toxin (PTX) blocks Gi-protein coupled receptor signaling, thereby preventing most chemokine-mediated cellular recruitment to the vaginal mucosa when applied intravaginally ^25^. Treatment with PTX ablated the protective effect of neomycin as mice treated with both pertussis toxin and neomycin prior to infection had significantly reduced vaginal ISG expression compared to PTX-untreated mice (Fig. 5a). This absence of ISG expression resulted in significantly higher levels of mucosal viral titers and an increase in disease pathology upon HSV-2 infection (Fig. 5b,c). These results suggested that cellular recruitment to the vaginal tissue is prerequisite to the enhanced ISG induction and protection against HSV-2 conferred by topical neomycin treatment.

**Figure 5:**
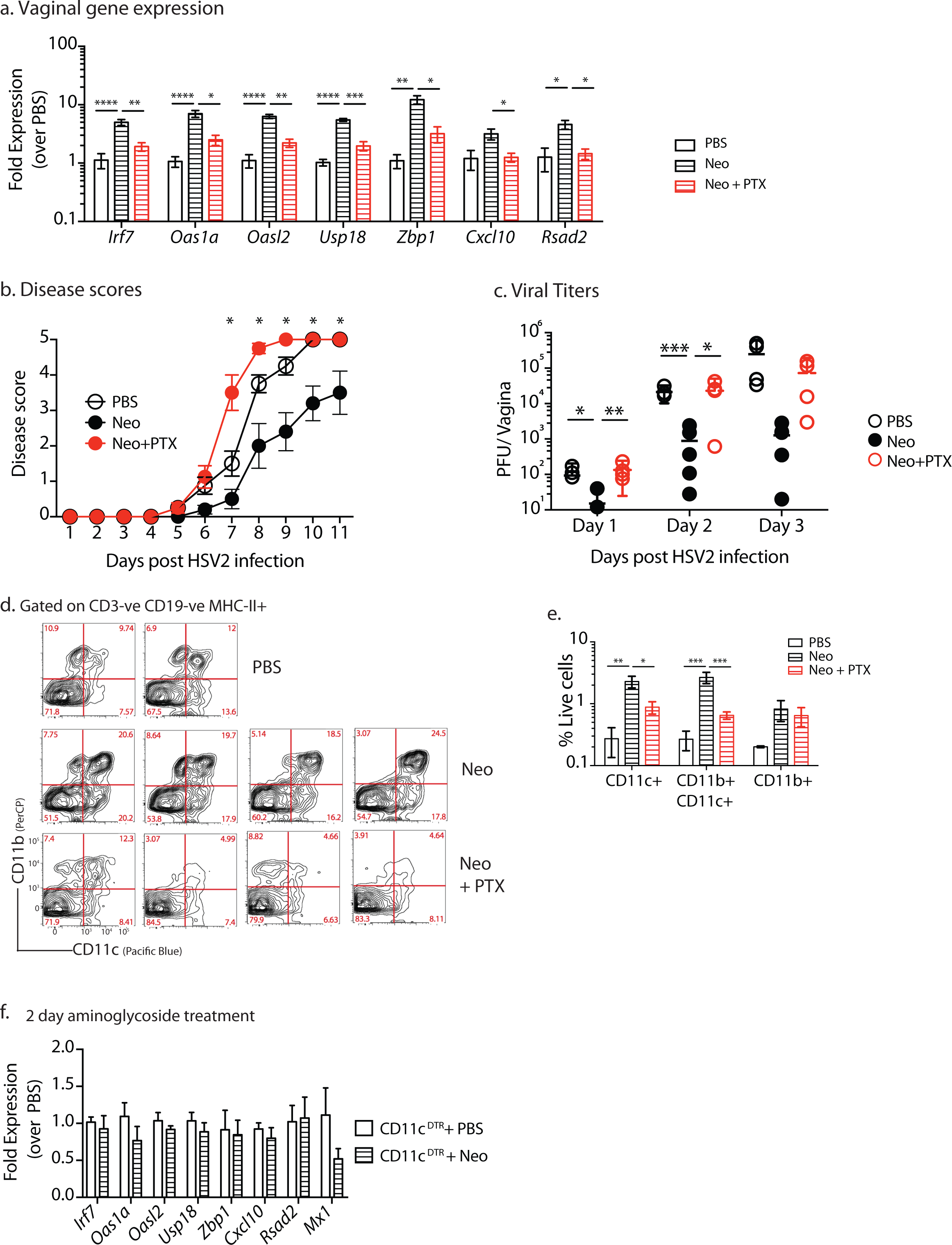
Recruited dendritic cells are required for ISG induction by neomycin. Mice treated subcutaneously with Depo-Provera were inoculated intravaginally with neomycin (1mg), PBS daily for 6 days (n = 4 mice per group) or neomycin (1mg) and pertussis toxin (PTX) (0μg) daily. After 1 week of treatment, vaginal gene expression was quantified (a). In an independent experiment, neomycin and PTX treated mice were infected with HSV-2 and disease score monitored daily (b) and vaginal viral titers measured (c). Vaginal dendritic cell populations were analyzed via flow cytometry, plots are gated on CD19-,CD3-, Gr1-, MHCII+ (d) and quantified in (e) (gating schema in Supplementary Fig. 14). Depo-treated CD11cDTR mice were treated with 125ng diphtheria toxin to deplete dendritic cells, treated intravaginally with 1mg neomycin for two days (n=8) and vaginal gene expression measured via qPCR (f). Error bars represent SEM. Significance was calculated using either a 2-way ANOVA for (b e) or using unpaired t-tests, correcting for multiple comparisons. Exact p values for all comparisons are reported in Table S2.

To determine the specific cell types recruited to the vaginal mucosa upon neomycin treatment, we compared the cellular composition of the vaginal mucosa in mice treated with neomycin and those treated with neomycin and PTX. Monocytes (CD11b+), monocyte-derived DCs (CD11b+CD11c+) and classical DCs (CD11c+), were significantly increased upon neomycin treatment (Fig. 5d,e). However, PTX treatment of neomycin-inoculated mice resulted in significant blockade of CD11c+ and CD11b+CD11c+ cells (Fig. 5e), suggesting that one or both of these cell types might be responsible for the neomycin-induced ISGs in the vaginal mucosa.

To test this hypothesis, we isolated CD11c+ cells from the vaginal tissue of neomycin-treated and control mice and found significant increases in transcripts of ISGs (Supplementary Fig. 8). Likewise, these recruited CD11c+ cells were also found to express IFNb (Supplementary Fig. 9a). To determine whether DCs are required for the increase in ISG expression following neomycin application, we treated CD11c-DTR mice with diphtheria toxin to deplete DCs and then administered neomycin intravaginally. In the absence of DCs, neomycin treatment failed to increase vaginal ISG expression (Fig. 5f). Collectively, our results suggest that recruited DCs are necessary for aminoglycoside-mediated ISG expression.

### XCR1+DCs are recruited to the vaginal mucosa upon neomycin treatment and are required for ISG induction

Since DCs are required for ISG induction by neomycin, and both monocyte-derived and classical DCs are recruited to the vaginal mucosa, we wondered if TLR3 expression could be used to identify the specific DC subset involved. It is well known that TLR3 is expressed selectively by the cDC1 and not cDC2 (Refs. ^26,27^). Examining TLR3 expression across all DC subsets using the RNA sequencing datasets deposited in the Immunological Genome project ^28^ confirmed a single cDC subset with high levels of TLR3 expression – CD8α+ DCs (cDC1) from the thymus and spleen (Supplementary Fig. 10). In non-lymphoid tissue, these cells are characterized as CD103+ DCs and both subsets express high levels of TLR3 (Ref. ^29^). Recent studies have identified XCR1 as a defining cell surface marker for cDC1 subset ^30^. Thus, we measured the recruitment of XCR+CD103+ DCs in the vaginal mucosa. Neomycin treatment resulted in significant recruitment of CD103+ XCR1+DCs that were ablated upon treatment with pertussis toxin (Fig. 6a,b). To determine if this was the DC subtype responsible for ISG induction by aminoglycosides, we depleted mice of XCR1+ DCs ^31^ before intravaginal aminoglycoside treatment. Mice lacking XCR1+ DCs showed no induction of ISG expression upon aminoglycoside treatment (Fig. 6c). These data suggested that mucosal application of aminoglycosides induce ISGs in TLR3 and XCR+ DC-dependent manner to confer antiviral protection.

**Figure 6:**
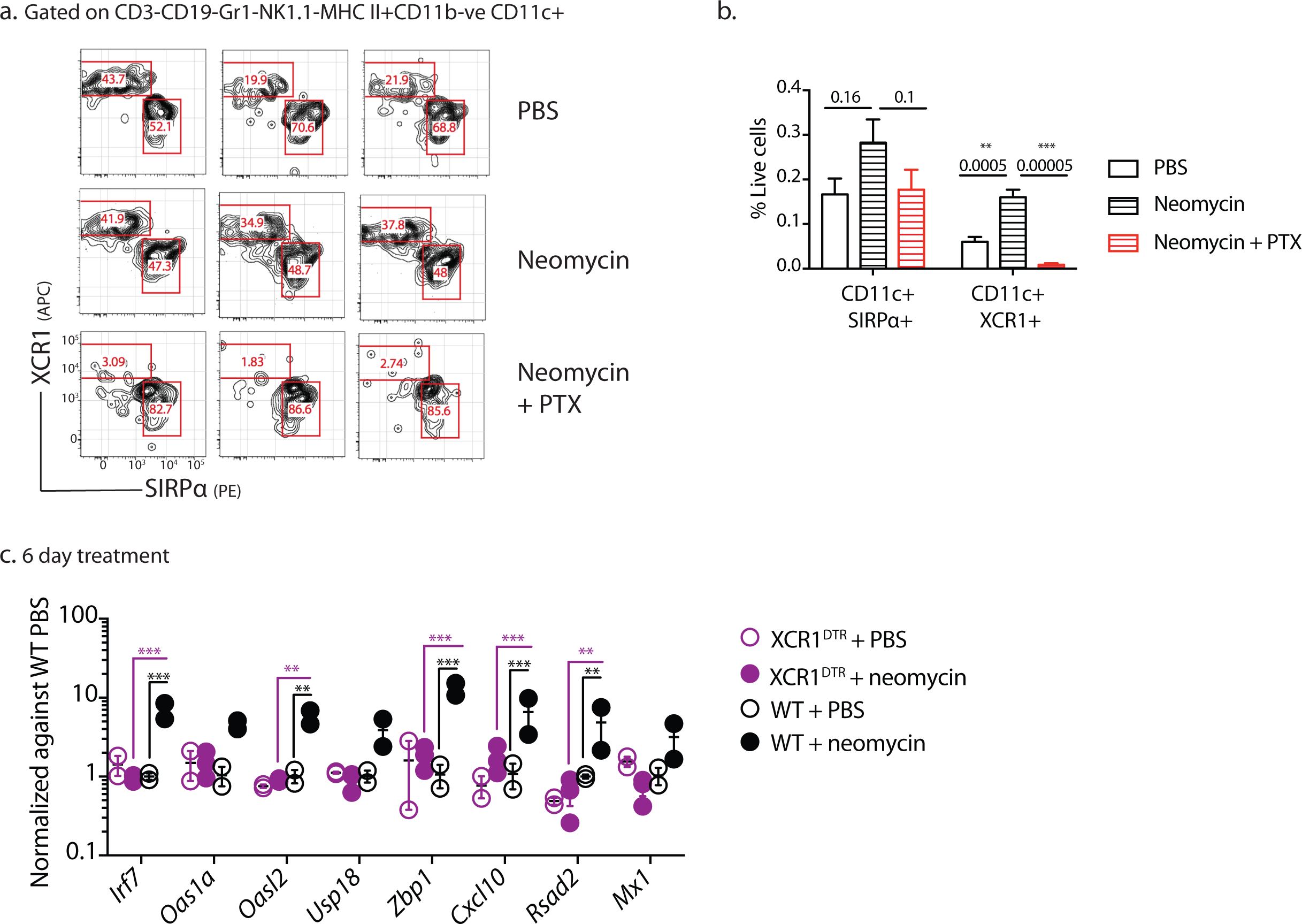
Recruited XCR1+ dendritic cells are required for ISG induction. Mice treated subcutaneously with Depo-Provera were inoculated intravaginally with neomycin (1mg), PBS daily for 1 week or neomycin (1mg) and pertussis toxin (PTX) (0μg) daily (n=3 mice per group). After 1 week of treatment, recruitment of XCR1 + vaginal DCs to the vaginal mucosa was measured. Plots are gated on CD19-, CD3-, Gr1-, MHCII+CD11c+ CD11b-live cells (a). Frequency of DCs quantified as a frequency of live cells (b) (n=4-6 mice per group). Depo-treated XCR1DTR mice were treated with 500ng diphtheria toxin to deplete XCR1+dendritic cells, treated intravaginally with 1mg neomycin for two days (n=2-5) and vaginal gene expression measured via qPCR (c). Exact p values for all comparisons are reported in Table S2.

Parenteral administration of aminoglycosides has known toxic side-effects including ototoxicity and nephrotoxicity ^32^. Aminoglycoside compounds can accumulate in the sensory hair cells of the inner ear causing caspase-mediated cell death which results in irreversible hearing loss ^33-35^. While nephrotoxicity can be reversible, significant necrosis can occur resulting in kidney dysfunction ^36,37^. However, we observed no toxicity in mice treated with intravaginal aminoglycosides. To determine if intravaginal aminoglycoside treatment resulted in mucosal toxicity, we conducted blinded histological analysis of aminoglycoside-treated and control vaginal tissues which found no histopathological differences between two groups (Supplementary Fig. 11a). Likewise, ex-vivo treatment of splenocytes with aminoglycosides induced robust ISG expression with little accompanying toxicity (Supplementary Fig. 11b-d. Similar to our in-vivo results, depletion of DCs resulted in significant loss of ISG induction (Supplementary Fig. 11e).

### Aminoglycoside enhances dsRNA stimulation of TLR3

Aminoglycosides acts by binding bacterial ribosomal RNA, but it also binds mitochondrial and mammalian ribosomal RNA^38-40^. Our data show that aminoglycoside induction of ISG requires TLR3, which is a sensor of dsRNA. Thus, we tested if aminoglycosides induce ISG expression by rendering host RNA more ‘visible’ to TLR3 in neighboring DCs. To test this, we first treated splenocytes with kasugamycin and washed the cells multiple times to remove extracellular aminoglycosides. Next, we incubated these kasugamycin-treated cells with splenic DCs that include XCR1+ DCs from WT and TLR3 knockout mice and measured DC-specific ISG expression. Incubation with kasugamycin-treated splenocytes was sufficient to increase ISG expression in WT but not TLR3^-/-^ DCs (Supplementary Fig. 12).

These data led us to hypothesize that aminoglycosides may bind to dsRNA and render them more potent for TLR3 activation. To test this, we treated splenocytes with a combination of aminoglycosides and dsRNA Poly I:C. We found that a 1:1000 ratio of Poly I:C to kasugamycin significantly induced ISG expression at greater levels than either compound alone (Supplementary Fig. 13). This enhancement was dependent on both TLR3 and TRIF signaling (Supplementary Fig. 13). Collectively, these data indicate that at high molar ratio, kasugamicin synergizes with dsRNA to stimulate TLR3.

## Discussion

Our data show an unexpected antiviral effect of aminoglycosides in mucosal tissues. Vaginal application of antibiotic cocktail conferred resistance against both DNA and RNA viruses. A single antibiotic, neomycin, was responsible for the protection and this protection extended to multiple members of the aminoglycoside family. Surprisingly, aminoglycoside-mediated antiviral protection occurred in germ free mice, indicating a microbiome-independent mechanism of resistance. Mucosal aminoglycoside treatment recruited XCR1+ dendritic cells, and induced ISGs expression via TLR3, TRIF and transcription factors IRF3 and IRF7. Finally, this antiviral protection could be extended to both the nasal mucosa and primary human monocytes.

Other antibiotic compounds have been previously reported to inhibit viral replication. Several screens of bioactive compounds for antiviral activity have identified antibiotic compounds including azithromycin and nanchangmycin ^41,42^. Azithromycin, a macrolide antibiotic, potently inhibits Zika virus replication in cell culture via an as yet unknown mechanism ^42^. Nanchangmycin was also originally identified in a screen for antiviral compounds effective against ZIKV. Pretreatment of cells with this antibiotic blocks entry of many flaviviruses including dengue virus and chikungunya virus by inhibiting clathrin-mediated endocytosis ^41^. Chloroquine, a member of the quinolone family of antimalarials has also demonstrated broad antiviral activity ^43-45^ and is thought to reduce viral replication by interfering with endosome acidification ^46^. These studies collectively highlight the unexpected antiviral functions of antibiotics, albeit through distinct downstream pathways. Our study identifies a class of antibiotics, aminoglycosides, which mediate their antiviral activity by increasing host expression of a broad range of ISGs, potentially reducing the opportunity for viruses to develop resistance. Of note, a previous study demonstrated the ability of anthracyclines (chemotherapy agents that are natural products of Streptomyces bacteria) to also induce ISG expression in cancer cells in a TLR3-dependent manner ^47^. Thus, bacterial products with nucleic acid binding capacity may have the ability to trigger TLR3 and possibly other PRRs.

How do aminoglycosides induce ISGs? Our results indicate that aminoglycoside treatment in the vaginal mucosa results in the increased expression of chemokines and is accompanied by recruitment of monocyte-derived DCs and cDCs. Blocking this recruitment through PTX treatment, or depletion of DCs, aborted the ISG response. With TLR3 as the key sensor required for ISGs and as only cDC1 expresses TLR3 (Ref. 29), we further determined that XCR1+ cDC1 were required for ISG induction in the vaginal mucosa.

Aminoglycosides could be taken up by these DCs via endocytosis ^48^ or through TRPV family of ion channels ^49^ which are also expressed on DCs ^50^. Aminoglycosides are known for inducing cytotoxicity in specific cell types such as the hair cells of the inner ear and kidney epithelial cells ^32,33,36^ but their effect on immune cells has not been well characterized. Aminoglycosides may also be taken up by the vaginal epithelial cells via the megalin receptor which has been shown to bind these compounds ^51,52^. While we observed no inflammation or cytotoxicity in aminoglycoside-treated vaginal tissues (Supplementary Fig. 11), we cannot rule out phagocytosis of nearby dead or dying cells by DCs. In support of this hypothesis, incubation of these DCs with kasugamycin-treated splenocytes was sufficient to induce ISG expression albeit at lower levels than direct aminoglycoside treatment (Supplementary Fig. 12). As XCR1+DCs are known for their cross-presentation of antigens associated with dead cells ^53,54^, it is conceivable that in the vaginal mucosa, phagocytosis of aminoglycoside-containing epithelial cells results in TLR3 activation due to the accumulation of aminoglycoside-bound RNA in the endosome.

The specific RNA bound by the aminoglycoside that triggers TLR3 remains unknown. The interaction of aminoglycosides with mitochondrial ribosomal RNA, which is closely related to bacterial ribosomal RNA, is well characterized and aminoglycosides have also been found to bind mitochondrial ribosomal RNA at multiple sites ^38,55^. It is possible that mitochondrial rRNA-bound aminoglycosides might activate TLR3 in XCR1+ DCs in the endosome. Our results show that aminoglycoside and Poly I:C can synergize to induce increased ISG expression (Supplementary Fig. 13). Future studies are needed to identify the manner in which aminoglycoside-bound RNA stimulate TLR3.

Our results demonstrate a surprising and broad antiviral effect of the aminoglycoside family of antibiotics, when applied to mucosal surfaces. However, we do not advocate for use of these compounds as antivirals, as aminoglycoside application is expected to cause local dysbiosis of commensal bacterial community. Understanding the precise mechanism by which aminoglycosides induce TLR3 stimulation will be useful for the future design of broad-acting antivirals.

## Materials and methods

### Mice

C57BL/6 (B6; Charles River Laboratories), B6(Cg)-Ifnar1tm1.2Ees/J (*Ifnar1*^-/-^), B6.FVB-Tg(Itgax-DTR/EGFP)57Lan/J (CD11cDTR), C57BL/6J-*Ticam1*^Lps2^/J (*Trif*^-/-^) mice, C57BL/6J-*Tmem173*^gt^/J (STING^-/-^) and B6.129S1-*Tlr7*^*tm1Flv*^/J (TLR7^-/-^), B6.129-*Ifnb1*^tm1Lky^/J (IFNβYFP reporter), B6;129S1-*Tlr3*^tm1Flv^/J (TLR3^-/-^) and B6129SF2/J controls were purchased from Jackson Laboratory unless otherwise specified and subsequently bred and housed at Yale University. *Irf3*^-/-^ *Irf7*^-/-^ (Ref. ^22^) (generous gift by Dr. T. Taniguchi, the University of Tokyo), *Mavs*^-/-^ (Ref. ^56^) (generous gift by Dr. Z. Chen, UTSW), B6.Cg-Xcr1^tm2(HBEGF/Venus)Ksho^ (XCR1-DTR) (Ref ^31^) are previously described. C57BL/6 mice carrying functional Mx1 alleles have were a generous gift by Dr. P. Staeheli ^57^ (University Medical Center Freiburg). Mice were maintained in our facility until the ages described. Germ-free Swiss Webster mice were maintained in flexible plastic gnotobiotic isolators with a 12 hr light/dark cycle and provided a standard, autoclaved mouse chow (5K67 LabDiet, Purina) ad libitum and autoclaved water for the duration of the experiment. Germ-free status was monitored by culture-based (aerobic and anaerobic culturing) and culture-independent methods (16S-targeted PCR). All procedures used in this study complied with federal and institutional policies of the Yale Animal Care and Use committee.

### Viruses

Wild-type HSV-2 (strain 186syn+) was a kind gift from Dr. David Knipe (Harvard Medical School, Boston, MA). HSV strain was maintained and propagated in Vero cells. Highly virulent variant of A/PR8 (Ref. ^19^) was a generous gift from Dr. P. Staeheli (University Medical Center Freiburg). Influenza virus strain A/PR/8/34 (H1N1) was propagated as previously described ^18^. ZIKV Cambodian FSS13025 strain was obtained from the World Reference Center for Emerging Viruses and Arboviruses at University of Texas Medical Branch, Galveston.

### Mouse infections and antibiotic treatment

C57BL/6 mice between six and seventeen weeks of age were subcutaneously injected with 2 milligrams of Depo-Provera in the neck scruff. Five days after Depo treatment, the mice were vaginally swabbed with a calcium alginate swab (Puritan, Maine) to remove vaginal mucus. The swab was wetted in sterile PBS and blotted on sterile paper to get rid of excess liquid before being used. Ten to 15 microliters of antibiotic or PBS was delivered into the vaginal cavity using a pipette tip. After 2-6 days of daily antibiotic treatment, mice were infected intravaginally with 2 × 10^3^ to 5 × 10^3^ PFU HSV-2, 10^6^ PFU HSV-1 or 2.5 × 10^5^ PFU ZIKV. Infections were carried out 24 hours after the last antibiotic treatment. At the time of infection, mice were weighed and subsequently examined at the same time each day to minimize fluctuations in weight due to circadian rhythms. Vaginal viral titers were collected by swabbing mice and flushing the vaginal cavity with 50 microliters of PBS. Mice were monitored daily for signs of inflammatory pathology scored as follows: (0) no inflammation, (1) genital inflammation, (2) genital lesions and hair loss, (3) hunched posture and ruffled fur, (4) hind-limb paralysis and (5) premoribund. Mice were euthanized before reaching a moribund state. Intranasal influenza infections were conducted as described previously ^18^. Briefly, mice were anesthetized using a mixture of ketamine and xylazine injected i.p. and infected with 500 PFU highly virulent A/PR8 influenza strain. All mouse experiments were carried out with a minimum of 3-4 mice per treatment group. With the exception of the germ-free mouse infections and experiments with XCR1^DTR^ mice, all other experiments were independently repeated at least once. These experiments were only conducted once with a total n=5. All animal procedures were performed in compliance with Yale Institutional Animal Care and Use Committee protocols.

### Antibiotic treatment

The antibiotic cocktail consisted of 0.5mgs Ampicillin, Vancomycin and Neomycin and 0.01 milligram metronidazole in a 15μl volume. For subsequent experiments using neomycin alone, mice were treated with 1 milligram in a volume of 10μΚ For intranasal treatment, mice were anesthetized by injecting a mixture of ketamine and xylazine intraperitoneally and 20μL of antibiotic administered dropwise into the nasal cavity using a pipet tip. All antibiotics were obtained from Sigma-Aldrich (Darmstadt, Germany).

### Microarray analysis

Mice were treated daily with neomycin (1mg/day) or PBS for six days and then sacrificed and vaginal tissue harvested. RNA was extracted from the vaginal tissues using an RNeasy extraction kit (Qiagen,CA) and hybridized onto MouseWG-6 v2 Expression arrays (Illumina,CA) at the Yale Center for Genome Analysis. Microarray data was visualized in R using ggplot2.

### Gene expression analysis

The following primers were used for mouse ISG expression *Hprt* (FP: GTTGGATACAGGCCAGACTTTGTTG, RP: GAGGGTAGGCTGGCCTATTGGCT); *Zbp1* (FP: TTGCCAATTCAAACGCCATC, RP: CACTTGTTGGAGCAAGGACT); *Cxcl10* (FP: AGAATGAGGGCCATAGGGAA, RP: CGTGGCAATGATCTCAACAC); *Usp18* (FP: CGTGCTTGAGAGGGTCATTTG, RP: GGTCCGGAGTCCACAACTTC); *Irf7* (FP: TGTAGACGGAGCAATGGCTGAAGT, RP: ATCCCTACGACCGAAATGCTTCCA); *Oas12* (FP: GGGAGGTCGTCATCAGCTTC, RP: CCCTTTTGCCCTCTCTGTGG); *Oas1a* (FP: ATTACCTCCTTCCCGACACC, RP: CAAACTCCACCTCCTGATGC); *Rsad2* (Viperin) (FP: AACAGGCTGGTTTGGAGAAG RP: TGCCATTGCTCACTATGCTC); *Mx1* (FP: CCAACTGGAATCCTCCTGGAA, RP: GCCGCACCTTCTCCTCATAG). The following primers were used for human ISG expression: *HPRT* (FP: TGGTCAGGCAGTATAATCCAAAG, RP: TTTCAAATCCAACAAAGTCTGGC); *OAS1* (FP: CTGAGAAGGCAGCTCACGAA, RP: TGTGCTGGGTCAGCAGAATC); *OASL* (FP: AAAAGAGAGGCCCATCATCCT, RP: CCTCTGCTCCACTGTCAAGT); *CXCL9* (FP: AGTGCAAGGAACCCCAGTAG RP: AGGGCTTGGGGCAAATTGTT); *CXCL10* (FP:CCACGTGTTGAGATCATTGCT RP: TGCATCGATTTTGCTCCCCT); *USP18* (FP:GGCTCCTGAGGCAAATCTGT RP: CAACCAGGCCATGAGGGTAG); *MX1* (FP: AGAGAAGGTGAGAAGCTGATCC; RP: TTCTTCCAGCTCCTTCTCCTG). qPCR reactions were run in triplicate. The triplicate Ct values, were averaged, normalized against housekeeping genes HPRT and then compared against biological controls (untreated mice) using the *Δ Δ* Ct method of comparison. Fold expression was calculated assuming a doubling efficiency (2) per cycle. (Fold expression = 2^-ΔΔCt^)

### Flow Cytometry Analysis

Single cell populations were isolated from vaginal tissue as previously described ^58^. Briefly, vaginal tissue was minced into small pieces and digested first with Dispase II for 15 minutes and subsequently with a combination of DNase I (.045 mg/ml) and Collagenase (2 mg/ml) for 30 minutes. All digestions were carried out in a 37°C shaking water bath. All enzymes were obtained from Sigma-Aldrich. Cells were spun down and dead cells excluded using a live/dead stain (Molecular Probes, Thermo Fisher), then stained with appropriate antibodies, fixed with 1% Paraformaldehyde (Electron Microscopy Sciences, PA) and run through an LSR II (BD, NJ) equipped with a UV laser. FlowJo software (Tree Star, OR) was used to visualize and analyze cytometry data. Cells populations were analyzed as shown in the gating scheme (Supplementary Fig. 14).

### Antibodies

The following antibodies were used for this study, all purchased from Biolegend unless specified otherwise: CD45(104), CD3ε (145-2C11), CD11c (HL3, BD Biosciences), CD11b (M1/70), CD103 (2E7) XCR1 (ZET), SIRPα(P84), Gr1 (RB6-8C5, BD Biosciences), NK1.1(PK136), MHC class II I-A/I-E (M5/114.152), CD19 (6D5).

### Human Monocyte Isolation and Infection

Peripheral blood mononuclear cells were obtained from the New York Blood Bank Center (NY) and monocytes isolated using a negative selection kit (Stemcell Technologies, MA). Monocytes were treated with PolyI:C (Sigma) or aminoglycosides at 2μg/ml and 2 mg/ml respectively for 6 hours. Cells were washed once and infected with Influenza A virus strain A/PR/8/34 (H1N1) at a multiplicity of infection of 2. Cells were incubated with the virus for 1 hour in 0.1%BSA in PBS. Cells were then washed once and incubated in DMEM containing 10%FBS, Pen/Strep, Sodium pyruvate and Hepes. RNA was isolated using an RNeasy extraction kit (Qiagen, CA) and viral RNA quantified using primers against Influenza A PR8 Polymerase A (FP:CGGTCCAAATTCCTGCTGAT; RP: CATTGGGTTCCTTCCATCCA).

### Viral Titer

Vaginal washes were collected in 50 μl PBS and saved in 950 μl PBS supplemented with 1% FBS, 10 mg/ml glucose 0.5 μM MgCl_2_ and 0.9 μM CaCl_2_. Washes were added in serial dilutions to a confluent monolayer of Vero cells and plaques visualized via crystal violet staining. ZIKV genome was quantified via qRT-PCR as previously described ^20^. Briefly, cDNA prepared from vaginal washes was compared against a standard curve composed of purified ZIKV viral genomes. Primers against NS5 (F: GGCCACGAGTCTGTACCAAA; R: AGCTTCACTGCAGTCTTCC) were used to measure ZIKV RNA.

### In-vitro treatments

Single cell splenocytes were plated at the density of 5×10^6^ cells/ml isolated and treated with kasugamycin (2mg/ml or 0.02mg/ml) and PolyI:C (High Molecular Weight, Invivogen,CA) 2μg/ml or 0.02μg/ml) for 6 hours. Mixtures of PolyI:C and kasugamycin were incubated together for 30 minutes at 37°C before addition to cells. In Supplementary Fig. 13, splenocytes were treated for 12 hours, washed 5 times, stained with cell trace violet (Thermo Fisher, CA), and incubated with splenic DCs isolated using PanDC separation kit (Stemcell Technologies, MA). After 6 hour incubation, cell trace violet dim and negative cells were sorted using an FACSAria (BD, NJ) into RLT buffer for RNA extraction.

### Statistics

Gene expression data was analyzed using unpaired t-tests, assuming unequal standard deviation and correcting for multiple comparisons using the Holm-Sidak correction unless otherwise specified. Graphs depicting disease scores were analyzed using 2-way ANOVA with Holm-Sidak correction for multiple comparisons. Graphs depicting viral titers across a time course of infection were analyzed using 2-way ANOVA with no correction for multiple comparisons (Fisher's LSD) unless otherwise specified. Survival curves were analyzed using the log-rank (Mantel-Cox) test. P values not reported in the figures themselves can be found in Supplementary Table 2. All statistical analyses were performed in Graphpad Prism v7.0.

## Data Availability

The data discussed in this publication have been deposited in NCBI's Gene Expression Omnibus^59^ and are accessible through GEO Series accession number GSE94909 (https://www.ncbi.nlm.nih.gov/geo/query/acc.cgi?acc=GSE94909).

## Acknowledgments

We thank Dr. Yong Kong for his help analyzing the microarray data, and Huiping Dong for animal support. We thank Punya Biswal for help with visualizing the microarray data. This study was supported by funding from the NIH AI054359, R56AI125504, R01EB000487 and 1R21AI131284 (to AI). AI and AG are Investigator and Faculty Scholar of Howard Hughes Medical Institute. SG and MK are recipients of the James Hudson Brown - Alexander Brown Coxe Postdoctoral Fellowships at Yale University.

---

**Supplementary Figure 1:**
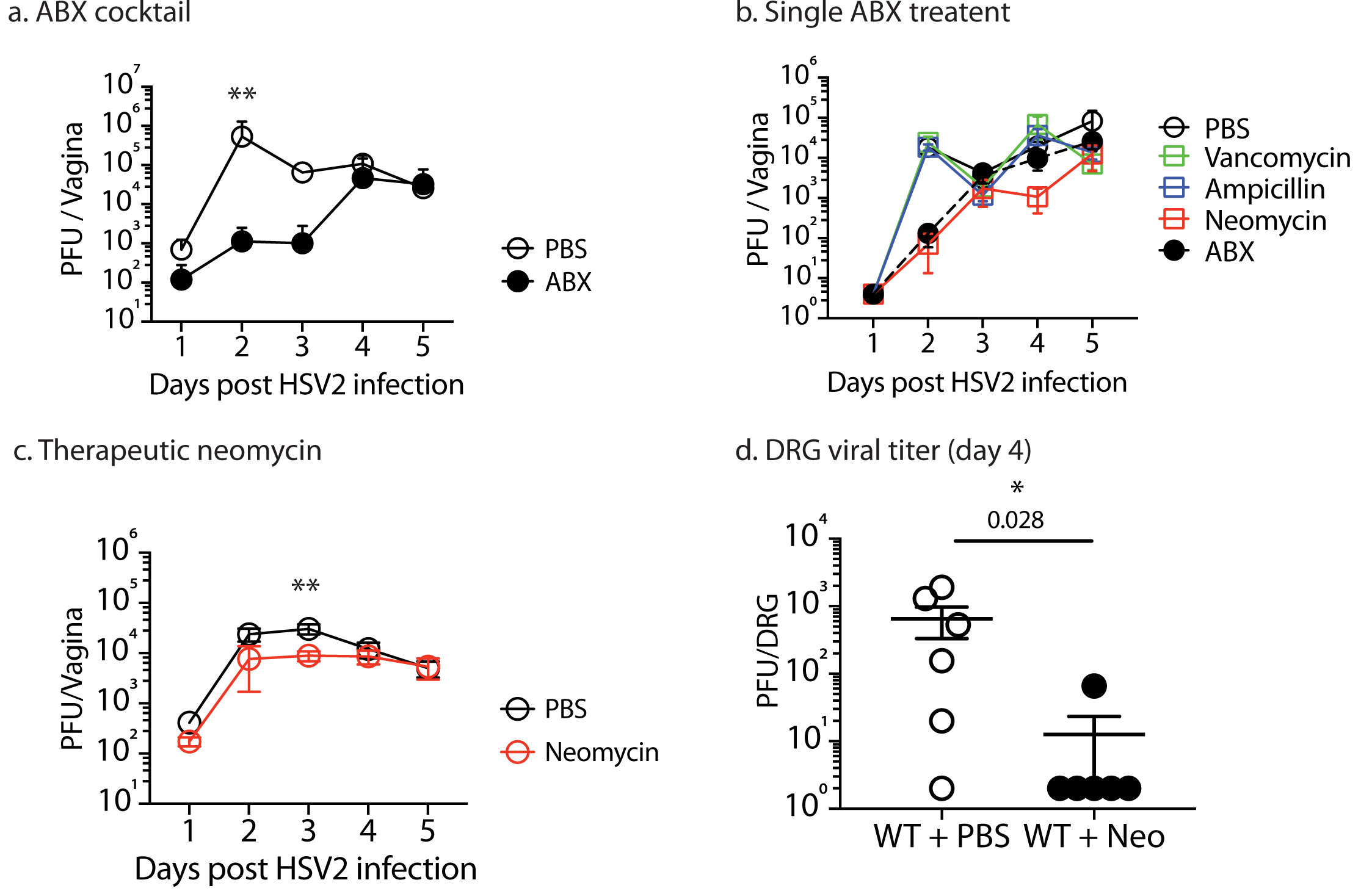
While vaginal viral replication of antibiotic-treated mice eventually reaches levels of PBS-treated controls, viral replication in dorsal root ganglion remain suppressed. Mice treated subcutaneously with Depo-Provera were inoculated intravaginally with the indicated aminoglycoside (1mg) or PBS daily for 1 week. After 1 week of treatment, mice were infected with HSV-2. Viral titers in the vagina were measured at indicated time points (a-c). Viral titers in the dorsal root ganglion were measured 4 days post infection (d). Error bars represent SEM and significance was calculated using unpaired t-tests. Exact p values are reported in Table S2.

**Supplementary Figure 2:**
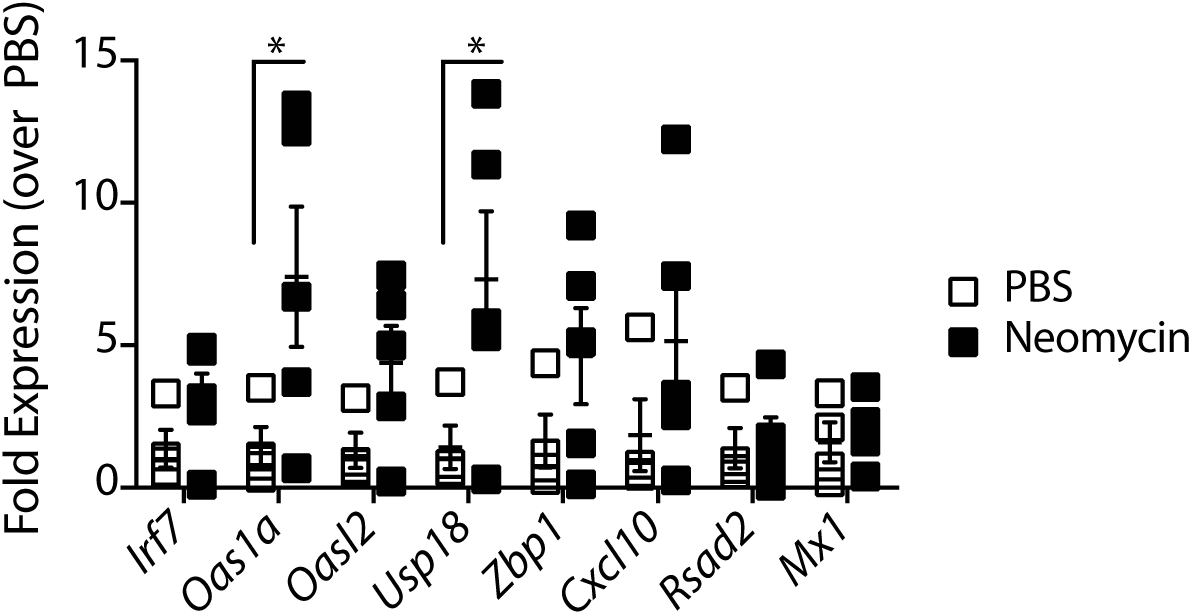
Neomycin treatment increases ISG expression in germ-free mice. Depo-treated outbred germ-free swiss-webster mice (n=4-5) were treated with 1mg neomycin or PBS daily for 6 days and vaginal gene expression analyzed. Error bars represent SEM and statistical significance was determined using a one-way ANOVA with * representing p values < 0.05. Exact p values are reported in Table S2.

**Supplementary Figure 3:**
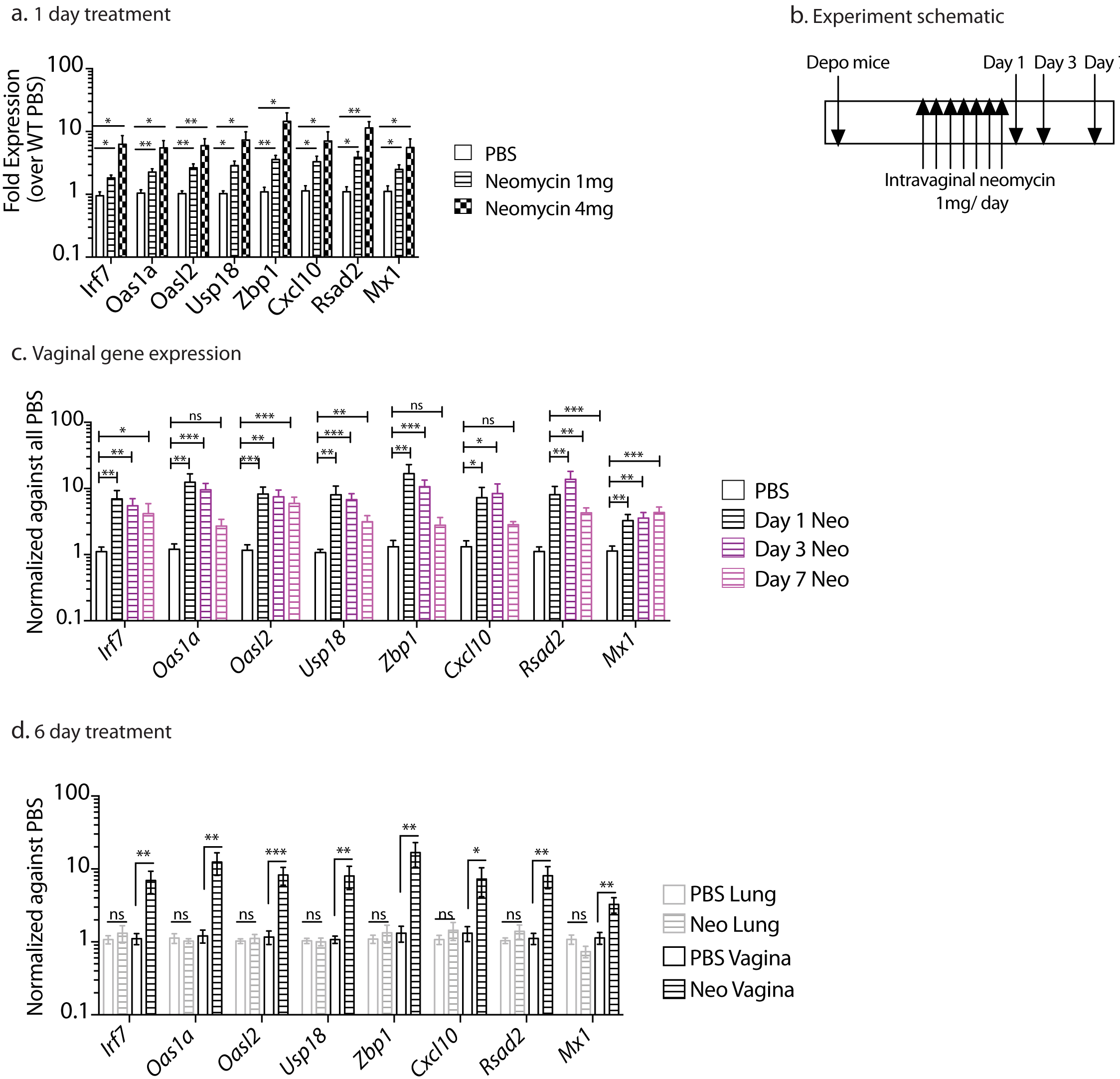
Kinetics of aminoglycoside-mediated ISG induction. Mice treated subcutaneously with Depo-Provera were inoculated intravaginally with neomycin (1mg or 4.5 mgs) or PBS for 1 day (n = 3-5 mice per group). 24 hours later, vaginal gene expression was quantified. Depo-treated mice were innoculated intravaginally with neomycin (1mg/day) or PBS daily for 6 days. Mice were sacrificed 1,3 and 7 days after the last neomycin treatment as shown in (b) and vaginal gene expression analyzed at each time point (c). Depo-treated mice were inncoluated with 1mg neomycin daily for 6 days and vaginal and lung ISG expression analyzed in mice 1 day post neomycin treatment (d). Error bars represent SEM and significance was calculated using unpaired t-tests, corrected for multiple comparison. Exact p values are reported in Table S2.

**Supplementary Figure 4:**
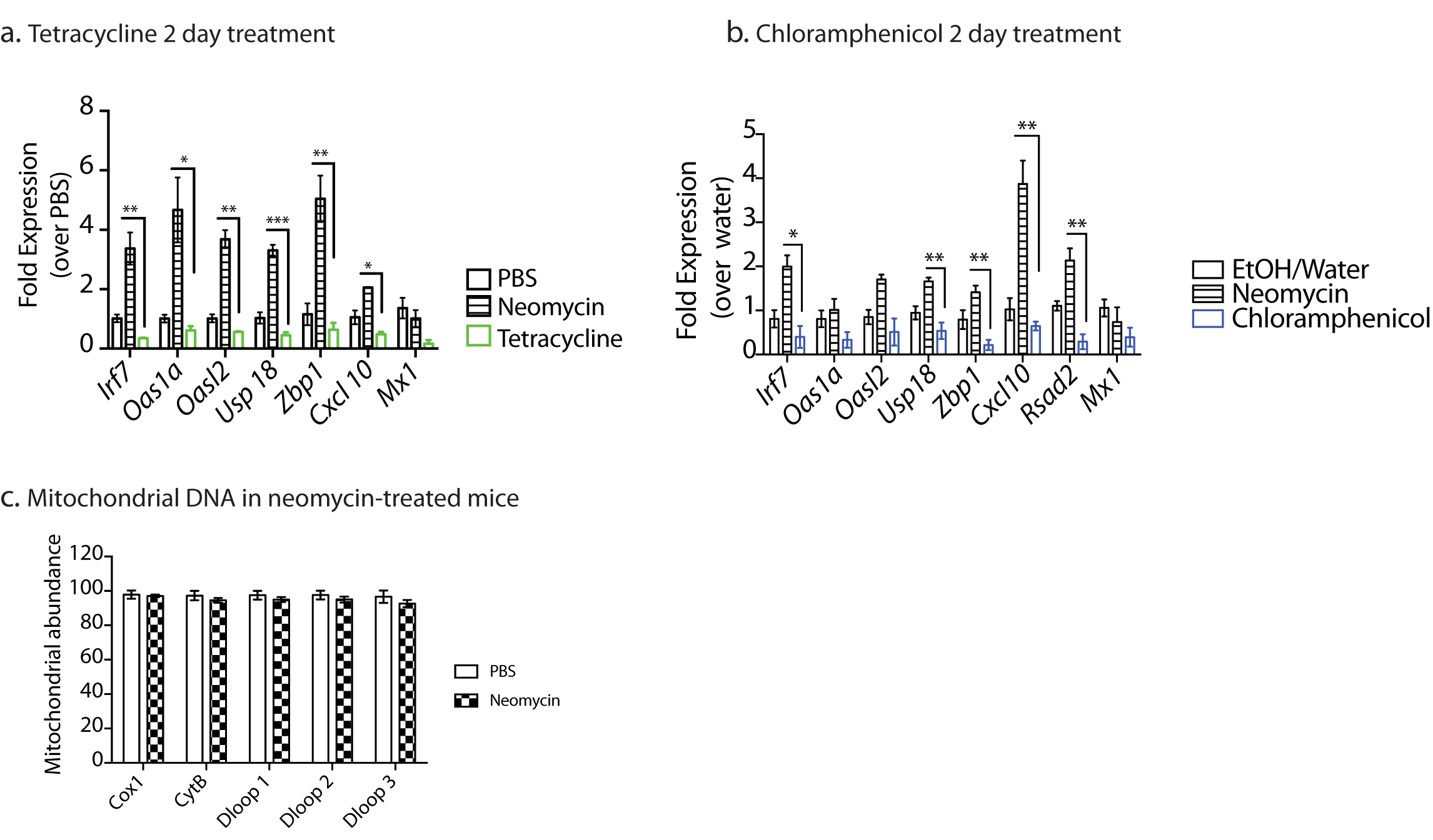
ISG induction is not a common property of ribosomal antibiotics. Mice were treated subcutaneously with Depo-Provera and five days after treatment were inoculated intravaginally with the indicated antibiotics (1mg/day) for 2 days (n=3 mice per group). Vaginal gene expression (a,b) and mitochondrial DNA abundance (c) was analyzed via qPCR. Significance was calculated using unpaired t-tests, correcting for multiple comparisons with * indicating p values <0.05. Exact p values for all comparisons are reported in Table S2.

**Supplementary Figure 5:**
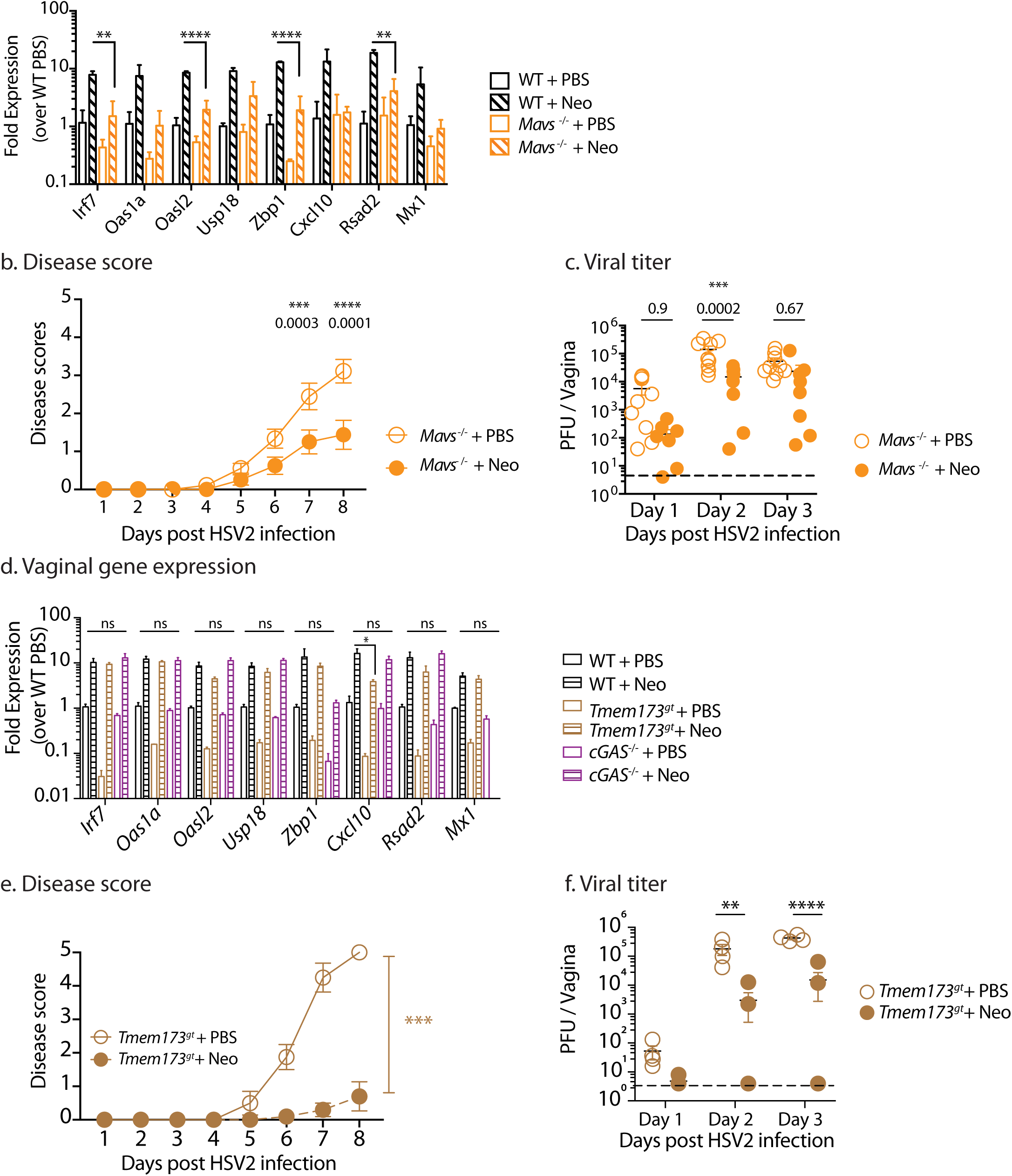
The role of RNA and DNA sensors in aminoglycoside-mediated antiviral protection. RNA sensor (*Mavs*^-/-^), DNA sensor (*Tmem173*^gt^, *cGAS*^-/-^) knockout mice and accompanying WT controls were treated subcutaneously with Depo-Provera and treated intravaginally with neomycin (1mg/day) or PBS for 6 days and vaginal gene expression measured via qPCR (a,d). Neomcyin-treated *Mavs*^-/-^ and *Tmem173*^*gt*^ mice were also infected with HSV-2, disease score monitored daily (b,e) and vaginal viral titers measured (c,f). Error bars represent SEM and significance was calculated using either unpaired t-tests (a,d) or 2-way ANOVA (b,c,e,f) correcting for multiple comparisons. Exact p values for all comparisons are reported in Table S2.

**Supplementary Figure 6:**
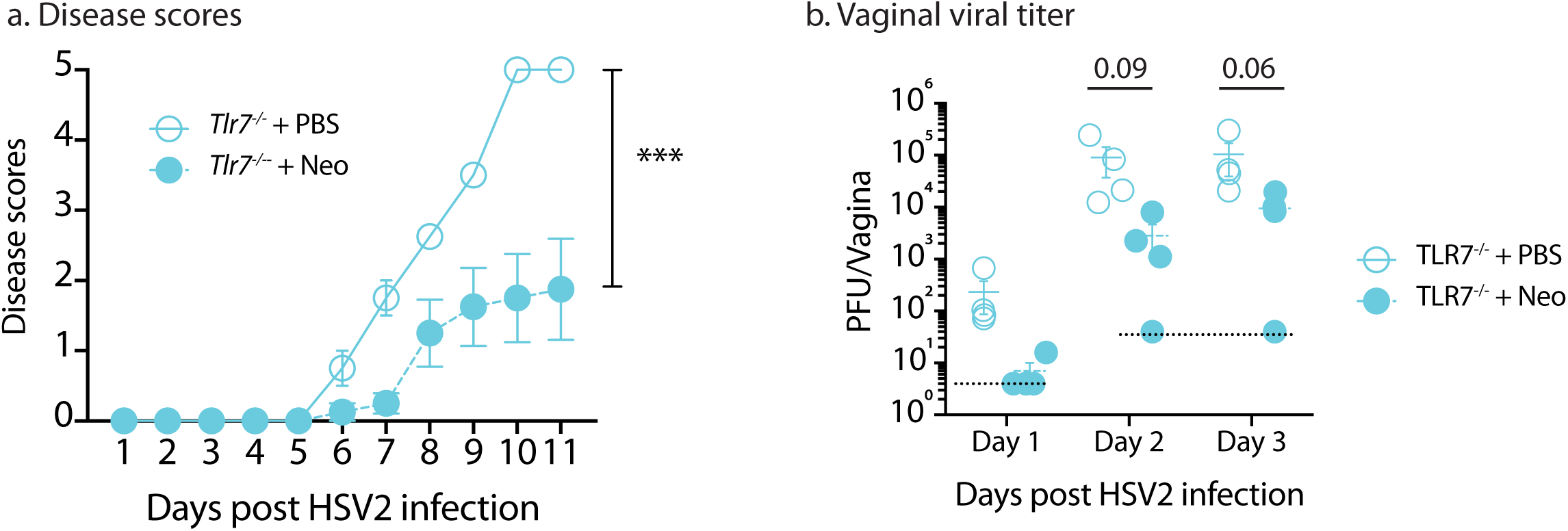
Aminoglycoside-mediated antiviral activity does not require TLR7 signaling. Depo-treated *Tlr7*^-/-^ mice were treated intravaginally with neomcyin (1mg/day) for 6 days and then infected with HSV-2 and disease scores monitored (a) and vaginal viral titer measured (b). Error bars represent SEM and statistical significance was calculated using 2-way ANOVA (a,b). Specific p values are reported in Table S2.

**Supplementary Figure 7:**
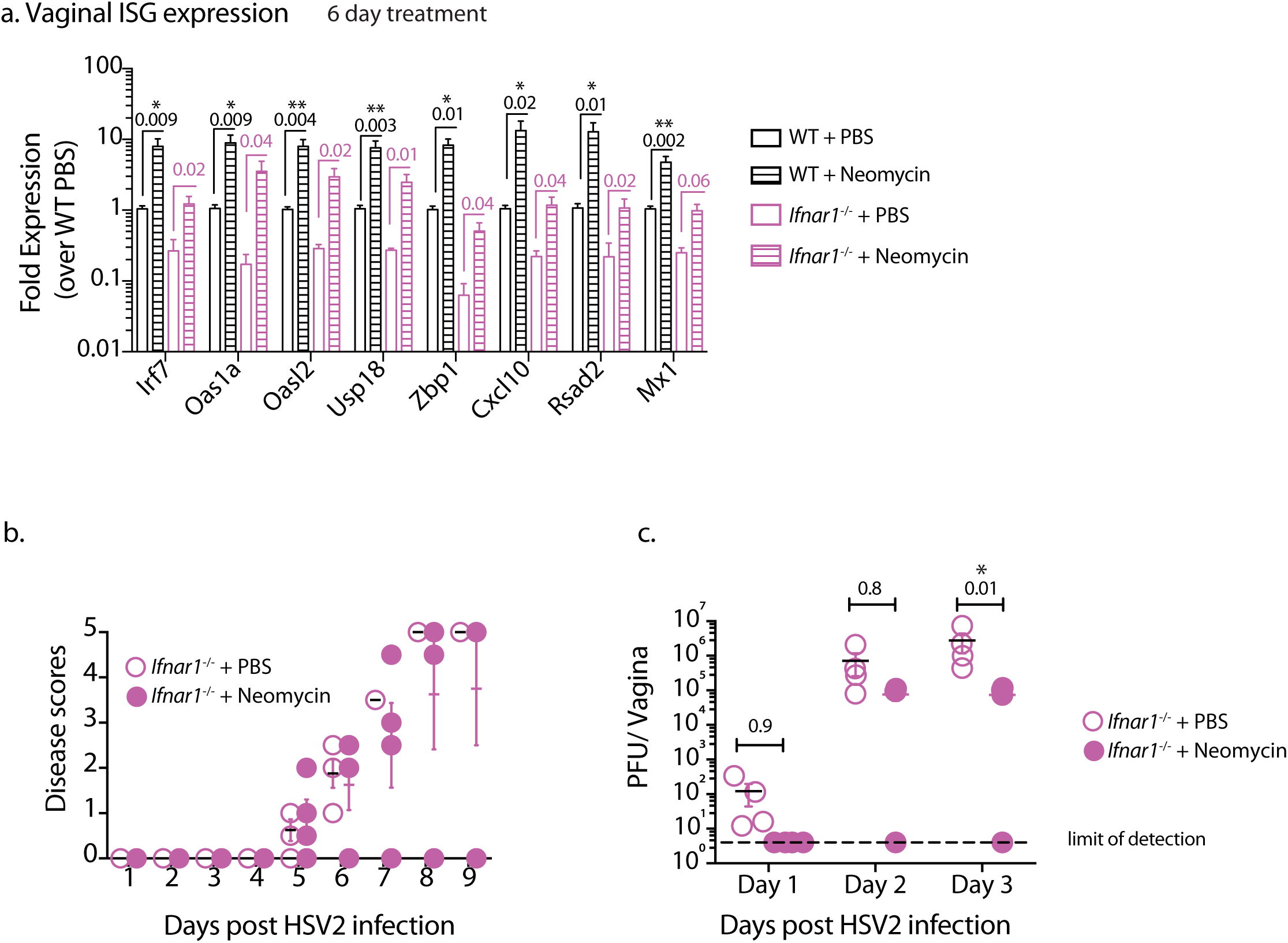
Role of IFNAR signaling in aminoglycoside-mediated antiviral protection. Type I IFN receptor knockout mice (*Ifnar1*^-/-^) and accompanying WT controls were treated subcutaneously with Depo-Provera and treated intravaginally with neomycin (1mg) or PBS daily for 6 days and vaginal gene expression measured via qPCR (a). In an independent experiment neomycin-treated *Ifnar1*^-/-^ mice and PBS-treated controls were also infected with HSV-2, disease score monitored daily (b) and vaginal viral titers measured (c). Error bars represent SEM and significance was calculated using either unpaired t-tests (a) or 2-way ANOVA (b,c) correcting for multiple comparisons. Exact p values for all comparisons are reported in Table S2.

**Supplementary Figure 8:**
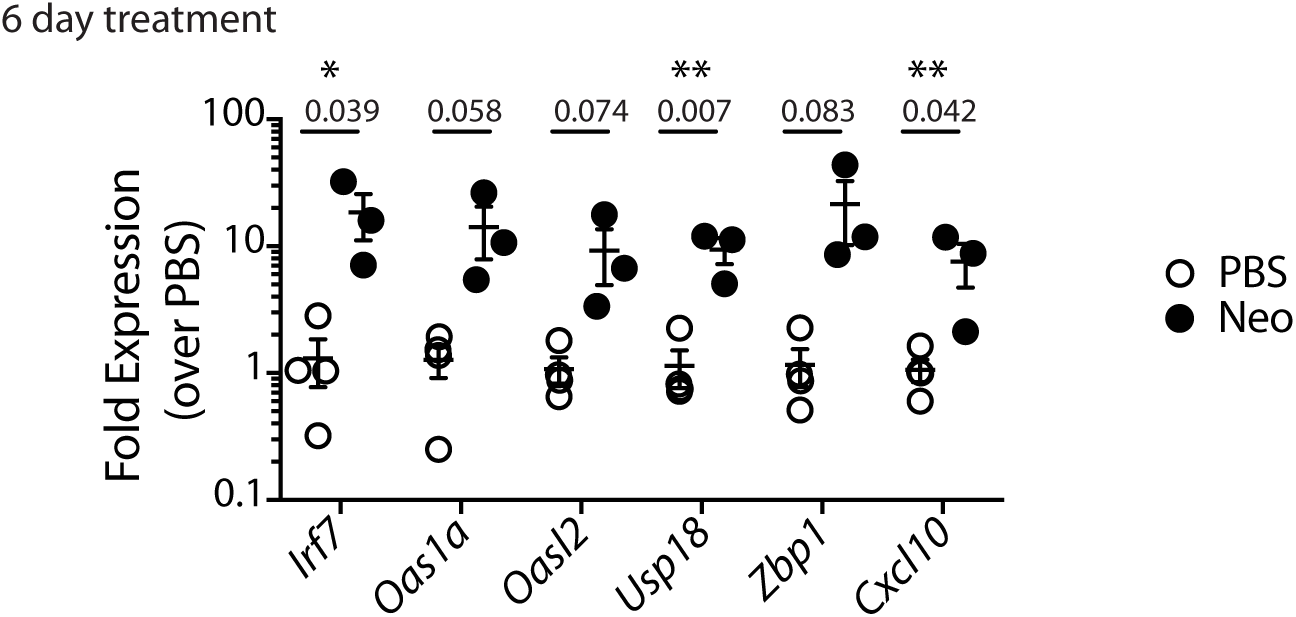
Neomycin treatment results in increased ISG expression in vaginal dendritic cells. Mice (n= 3-4) were treated subcutaneously with Depo-Provera and five days after treatment were inoculated intravaginally with neomycin (1mg) or equivalent volume of PBS daily for six days. Vaginal tissue was harvested, CD11c+ve cells isolated and gene expression analyzed. Significance was calculated using unpaired t-tests, correcting for multiple comparisons with * indicating p values <0.05.

**Supplementary Figure 9:**
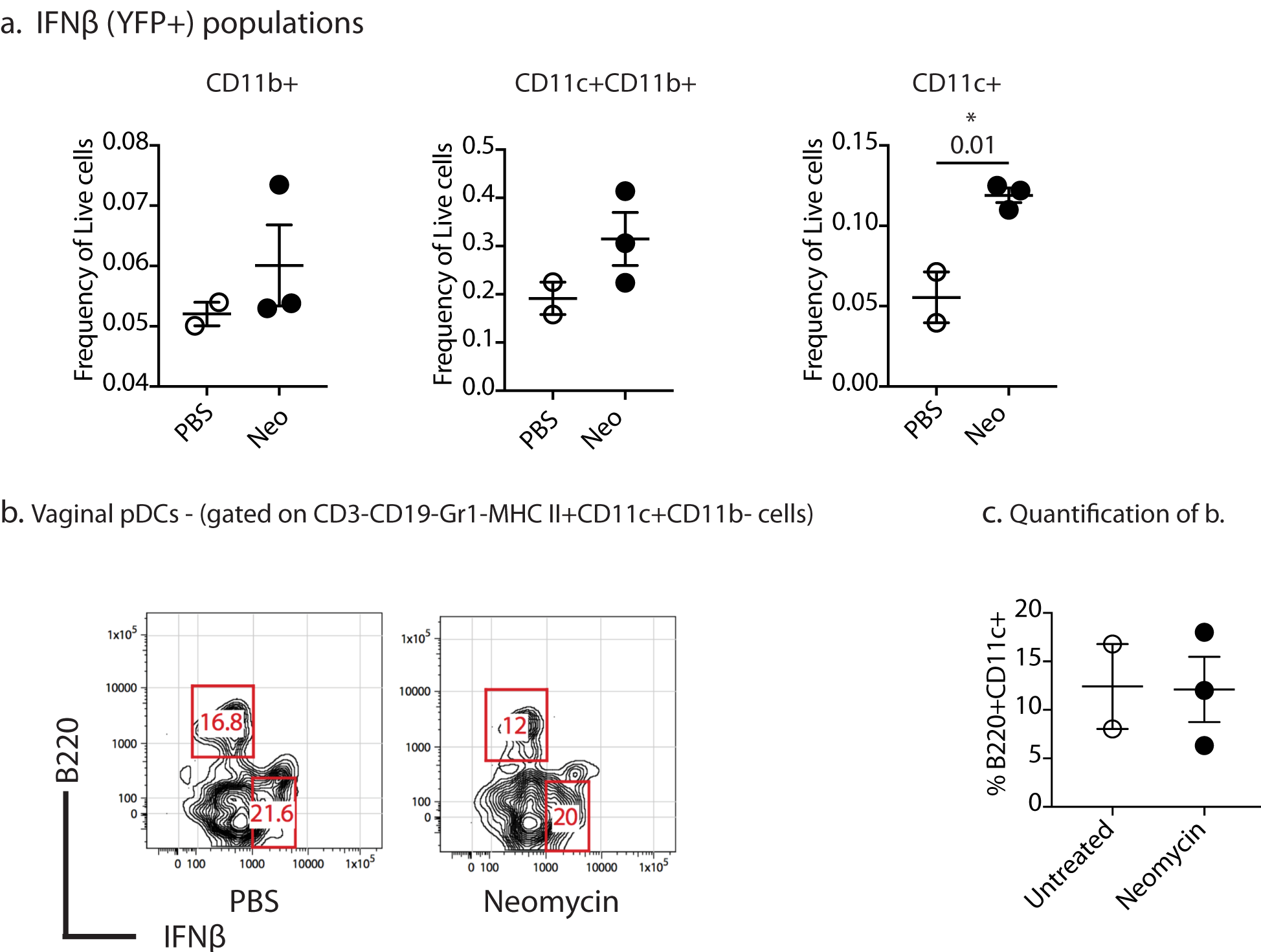
Neomycin treatment increases recruitment of IFNβ+ cDCs. Depo-treated IFNβ^YFP^ reporter mice were treated with 1mg neomycin or PBS daily for 6 days and vaginal dendritic cell populations analyzed. IFNβ populations were gated on CD3-CD19-NK1.1-Gr1-MHC II+ cells and quantified as a frequency of total live cells (a). Plasmacytoid DCs were gated as B220+CD11c+ cells and quantified (b,c). Error bars represent SEM and significance was calculated using an unpaired t-test.

**Supplementary Figure 10:**
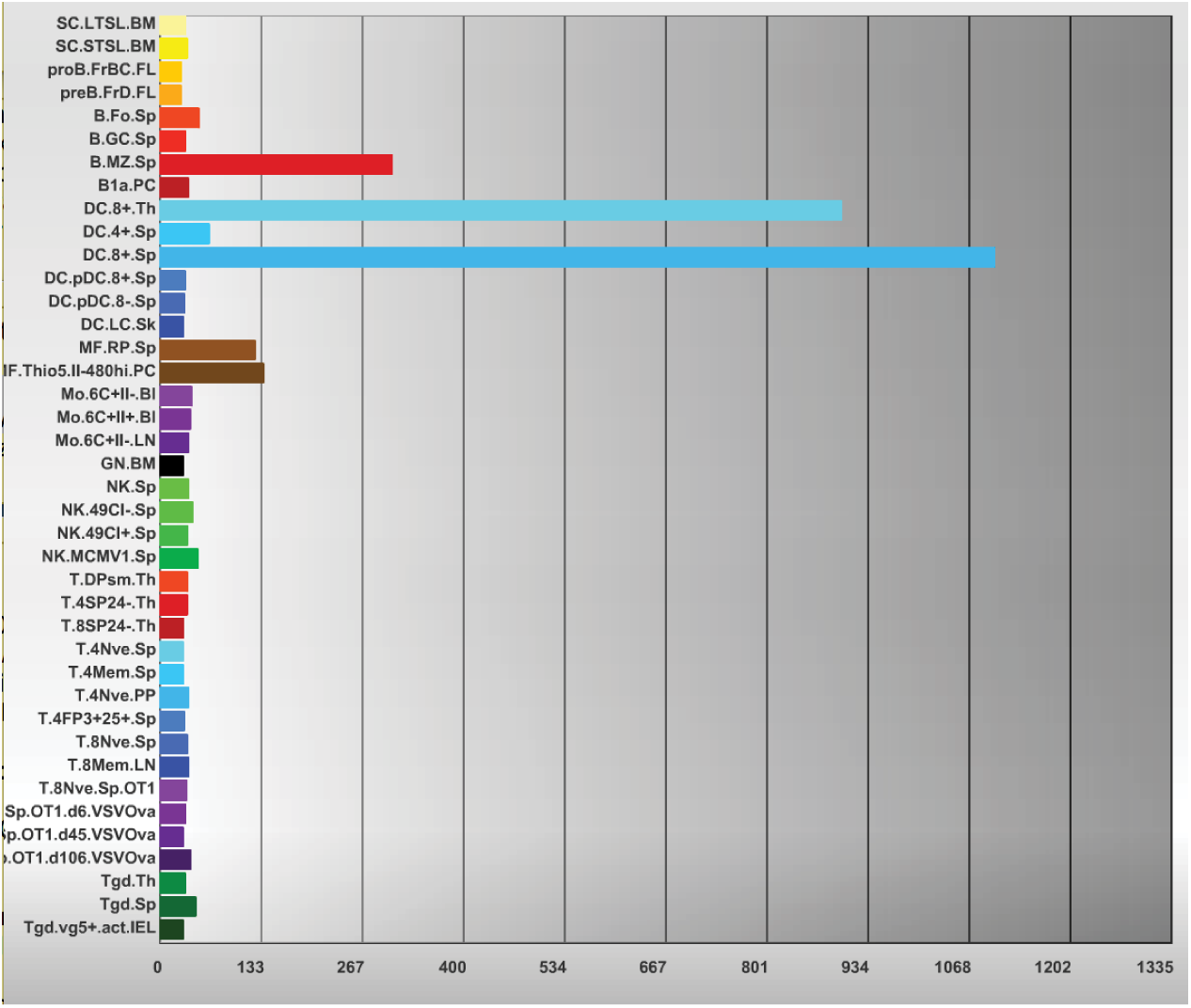
TLR3 expression across immune cell types obtained from the ImmGen database. The ImmGen data base (https://www.immgen.org/) was queried on 9/21/2017 for expression of TLR3 across all available dendritic cell gene expression datasets.

**Supplementary Figure 11:**
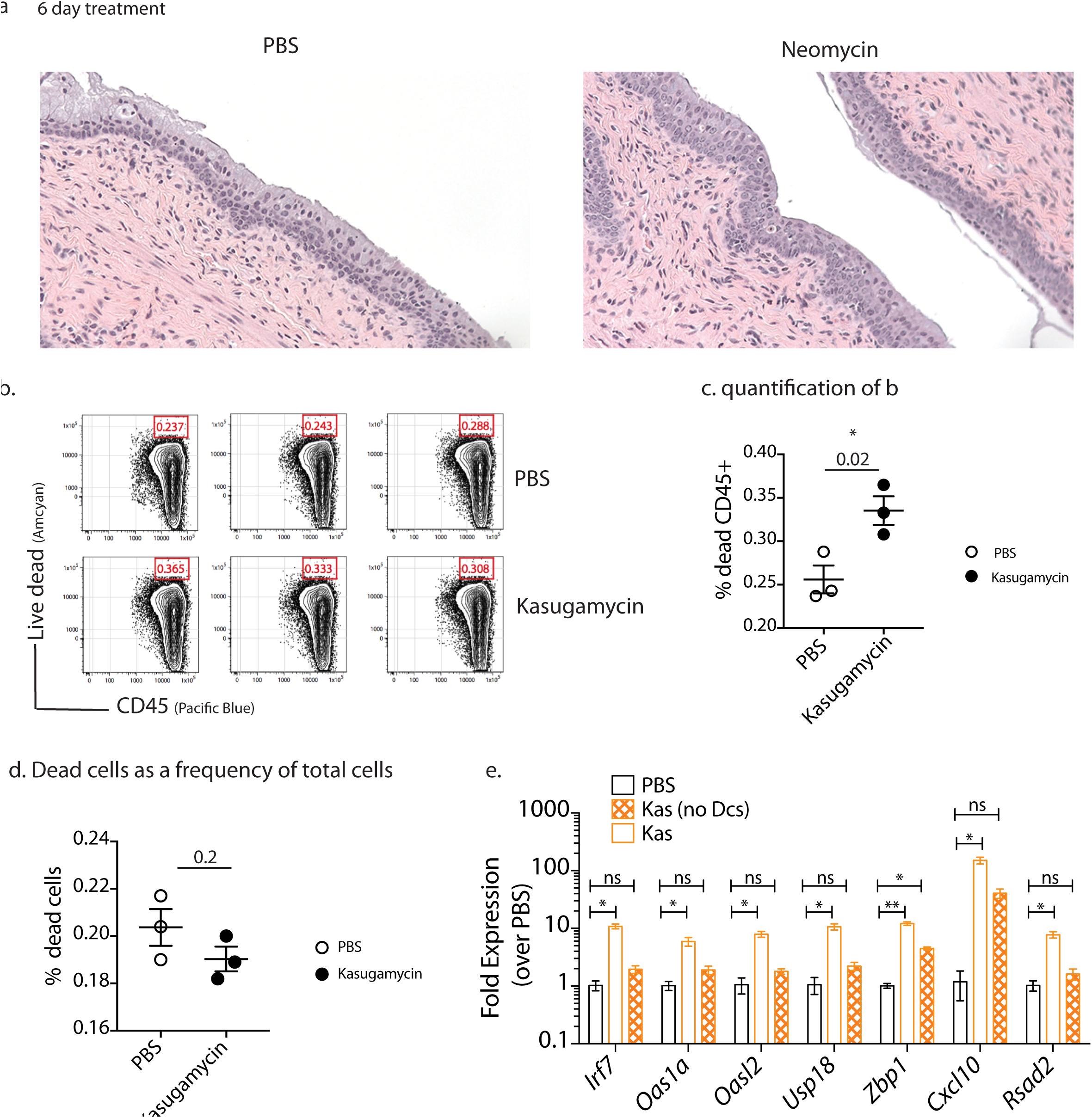
Intravaginal aminoglycoside treatment induces minimal inflammation and *in-vitro* aminoglycoside treatment induces minmal cell death. Mice treated subcutaneously with Depo-Provera were inoculated intravaginally with PBS or neomycin (1mg) for one week. Vaginal tissue from neomycin-treated and PBS control mice was fixed, embedded and stained with hematoxylin and eosin. Images are magnified 200X (a). Splenocytes were treated with 2mg/ml kasugamycin for 6 hours and dead cells were quantified using fixable live/dead stain as a frequency of total leukocytes (c) or total cells (d). In an independent experiment kasugamycin-treated splenocytes were depleted of DCs and ISG expression quantified (e). Error bars represent SEM and exact p values are reported in Table S2.

**Supplementary Figure 12:**
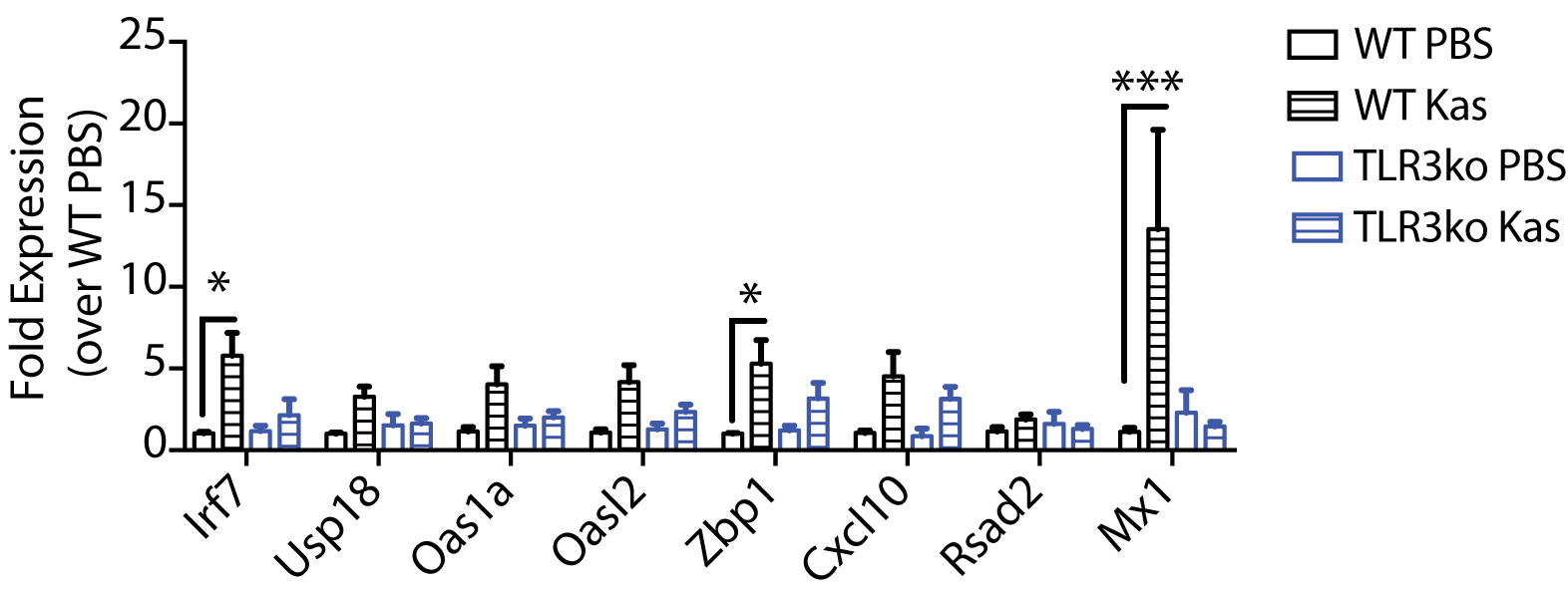
Kasugamycin-treated splenocytes induce ISG expression in DCs in a TLR3 dependent manner. Splenocytes were treated with2mg/ml kasugamycin for 12 hours, washed thoroughly, stained with cell trace violet and incubated with DCs isolated from WT or *Tlr3*^-/-^ splenocytes. After 6 hours of incubation, DCs (cell trace violet dim and negative cells) were sorted, RNA extracted and ISG expression quantified. Error bars represent SEM and specific p values are detailed in Table S2.

**Supplementary Figure 13:**
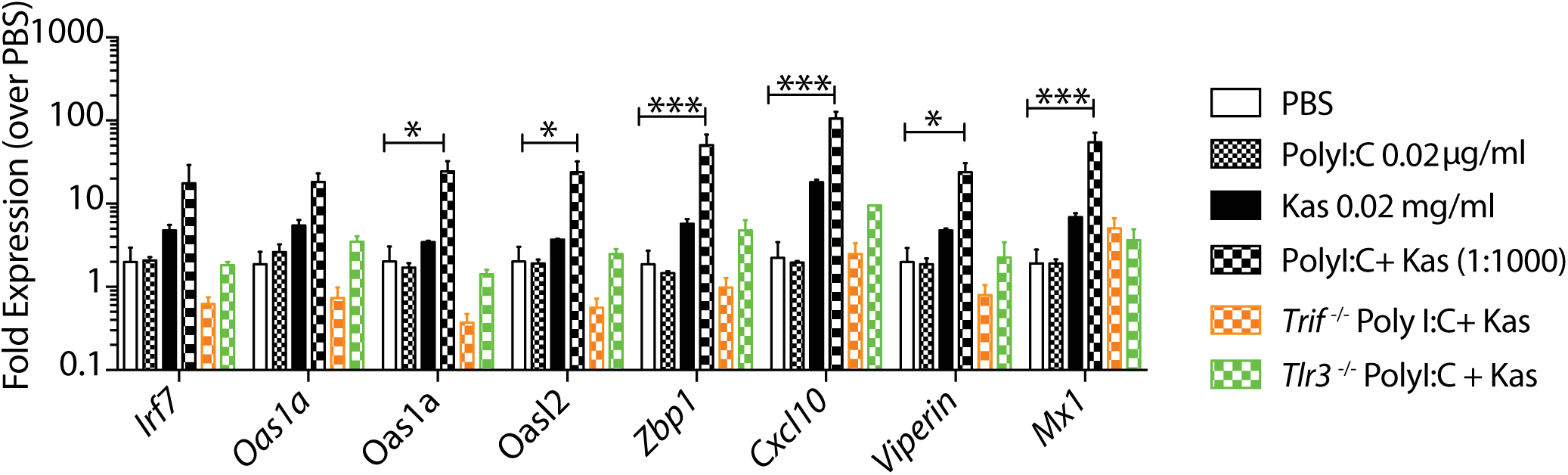
Kasugamycin can synergize with PolyI:C to amplify ISG expression. Splenocytes from WT, *Trif*^-/-^ or *Tlr3*^-/-^ mice were treated with 0.02μg/ml Poly I:C, 0.02mg/ml kasugamycin or the two together for 6 hours and ISG expression quantified. Error bars represent SEM and significance was calculated using a 2-way ANOVA. Specific p values are reported in Table S2.

**Supplementary Figure 14:**
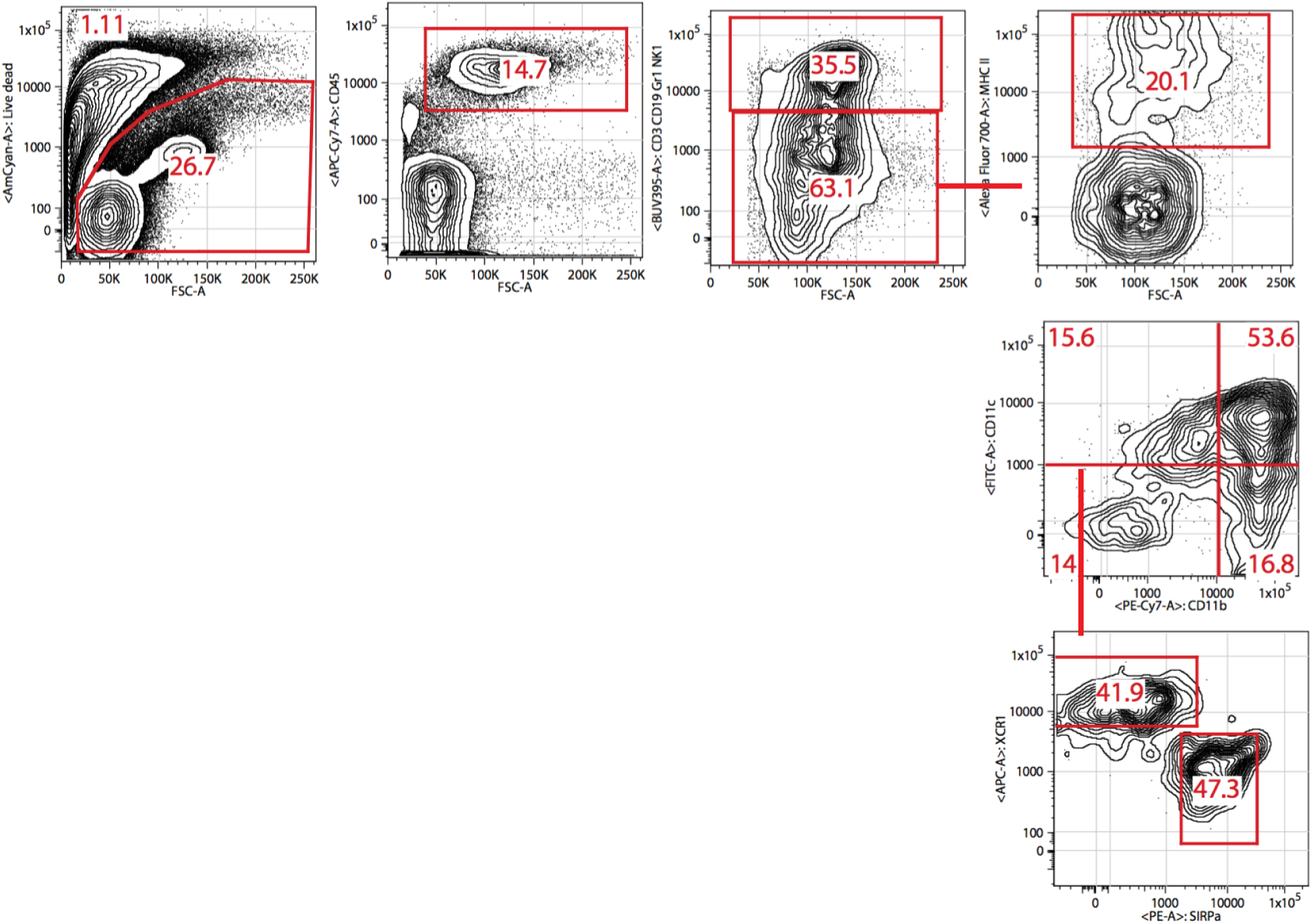
Flow cytometry gating schema. Vaginal single cell suspensions were gated on live cells, leukocytes (CD45+). MHC-II+ve cells were gated on CD3/CD19/G1-ve populations. CD11b-Cd11c+ cells were gated on MHC-II+ve cells and were further gated on XCR1 + SIRPα-ve cells.

